# Diversity and task-dependence of task representations in V1 during freely-moving decisions

**DOI:** 10.1101/2022.07.06.498845

**Authors:** Anqi Zhang, Anthony M. Zador

## Abstract

Neurons in primary visual cortex (area V1) are strongly driven by both sensory stimuli and non-sensory events. However, although the representation of sensory stimuli has been well characterized, much less is known about the representation of non-sensory events. Here, we characterize the specificity and organization of non-sensory representations in rat V1 during a freely-moving visual decision task. We find that single neurons encode diverse combinations of task features simultaneously and across task epochs. Despite heterogeneity at the level of single neuron response patterns, both visual and non-visual task variables could be reliably decoded from small neural populations (5-40 units) throughout a trial. Interestingly, in animals trained to make an auditory decision following passive observation of a visual stimulus, some but not all task features could also be decoded from V1 activity. Our results support the view that even in V1— the earliest stage of the cortical hierarchy—bottom-up sensory information is combined with top-down non-sensory information in a task-dependent manner.

## Introduction

The brain processes and transforms sensory inputs to generate appropriate motor outputs. How brain areas contribute to this goal is related to the features they can represent. In primary visual cortex, neural activity has historically been characterized in terms of stimulus parameters such as orientation, spatial frequency, temporal frequency, and direction of visual motion (Felleman and Van Essen 1991; Hubel and Wiesel 1959; Marques et al. 2018). By contrast, complex combinations of task-relevant and abstract features are often found in downstream areas in parietal and frontal cortices (Hanks et al. 2015; Krumin et al. 2018; Morcos and Harvey 2016; Raposo et al. 2014; Scott et al. 2017). Although it has been long been recognized that sensory cortices are not driven solely by bottom-up sensory inputs—the first single unit recordings reported attentional modulation of auditory responses in the cat (Hubel et al. 1959)—there has recently been growing recognition of the importance of non-sensory responses in primary visual cortex (V1), such as those related to locomotion, arousal, and body movements (Musall et al. 2019; Niell and Stryker 2010; Vinck et al. 2015).

The role of non-sensory responses in primary sensory cortices remains an open question. Although sensory representations in primary sensory cortices are important for perceptual decisions, the magnitude of stimulus-evoked activity in sensory cortices is frequently overshadowed by the magnitude of activity due to task-condition, movement or outcome (Musall et al. 2019; Niell and Stryker 2010; Orsolic et al. 2021; Otazu et al. 2009; Shuler and Bear 2006; Stringer et al. 2019). Non-sensory signals both modulate and appear independently of sensory-related activity in primary visual and auditory cortices (Guitchounts et al. 2020; Jaramillo and Zador 2011; Keller et al. 2012; Musall et al. 2019; Niell and Stryker 2010; Shuler and Bear 2006; Steinmetz et al. 2019; Stringer et al. 2019). In V1, non-sensory representations may support some visual computations, such as computing visual expectations during virtual reality locomotion or navigation, and in these cases are coherent with relevant sensory representations (Fiser et al. 2016; Keller et al. 2012; Poort et al. 2015; Saleem et al. 2018). However, non-sensory driven activity has also been observed when such computations are not necessary, and can both correlate with and occur independently of task variables (Musall et al. 2019). We set out to understand how task-related non-sensory activity is organized and how it relates to sensory encoding and task demands.

Here we used extracellular methods to record responses from single neurons in area V1 of freely moving rats performing a visual discrimination task. We find that neurons in this area encode both sensory and non-sensory task variables. In control animals trained to perform a similar but non-visual task, the encoding of sensory stimuli was similar, but the fidelity with which some non-sensory task variables were encoded was markedly diminished. Our results demonstrate that even in V1—the earliest stage of the cortical hierarchy—bottom-up sensory information is combined with top-down information in a task-dependent manner.

## Results

In what follows, we first describe a visual spatial discrimination paradigm for freely moving rats, along with software methods to constrain the animal’s viewing position and angle. Then, we characterize visual and nonvisual representations in V1 single neuron activity recorded using tetrode microdrives. We analyze this activity for representations of task parameters such as stimulus, choice, movement parameters, and outcome. We then investigate whether neurons are specialized for encoding single task features, or are influenced by combinations of task features within and across task epochs. Conversely, we ask to what extent these task features can be read out from neural populations at various points in the task. Finally, we compare V1 response profiles during the visual spatial discrimination to those during an analogous but visually-independent task.

### “Cloud of dots” visual discrimination task

To probe the patterns of representations in primary visual cortex during a freely moving visually guided behavior, we first designed a fixed-time visual discrimination task for freely moving rats. Rats were placed into a behavior chamber containing three nosepokes (Uchida and Mainen 2003, Otazu et al. 2009). Rats self-initiated trials by poking into the center stimulus viewing port, and were presented with a 500ms-long visual stimulus of distributed flickering dots (Figure 1a). They were asked to judge the region of higher dot density (top versus bottom) presented in the stimulus and reported their decision by poking into one of the side nosepokes after delivery of a decision tone signaling the beginning of the decision period. Correct choices earned a small water reward, while incorrect choices earned a punishment tone and time-out.

**Figure 1.**
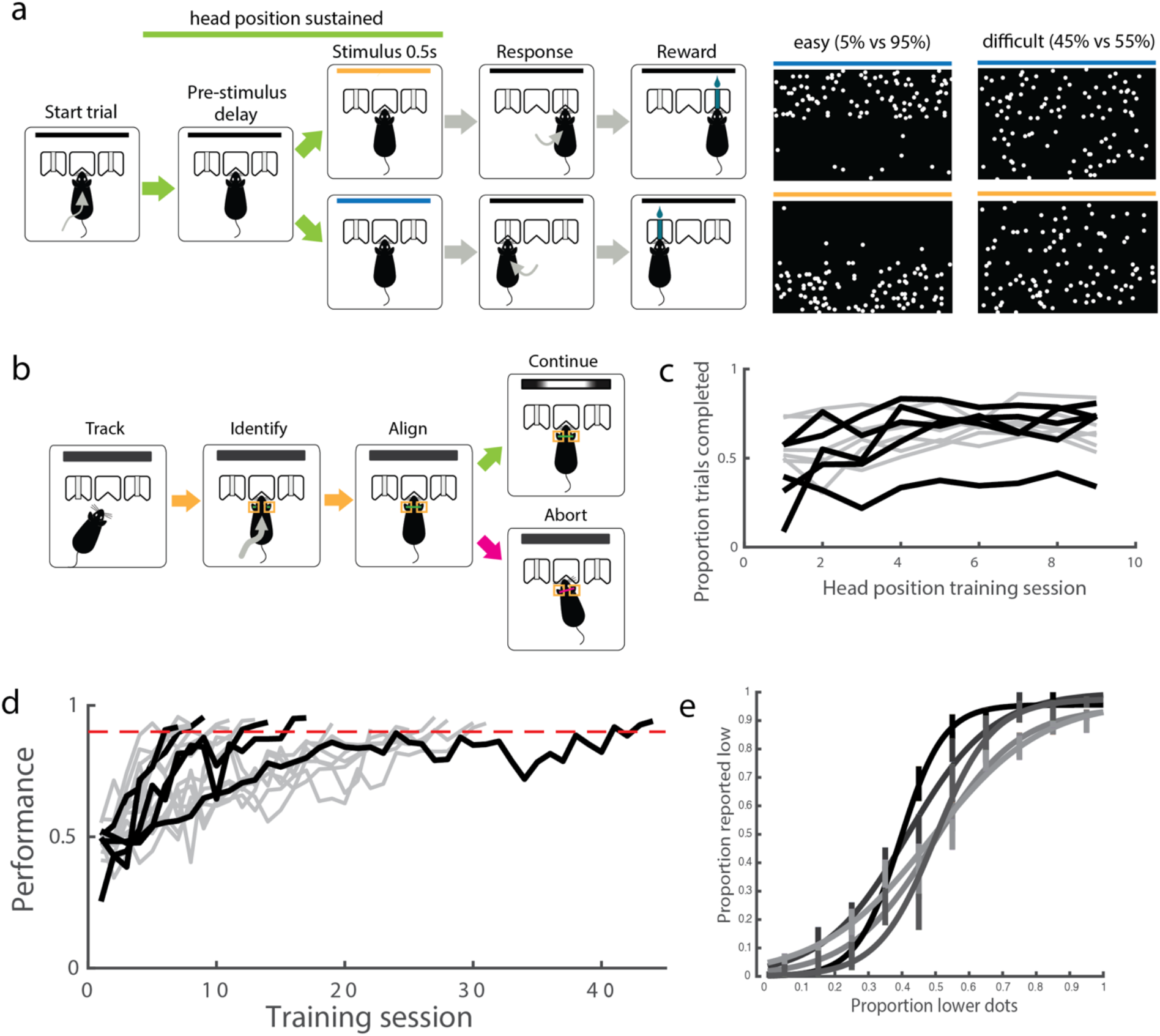
Rats reliably learn a “cloud of dots” visual discrimination task. a. Task design, with example stimulus frames for upper hemifield (top) and lower hemifield (bottom) trials (left: easy, right: difficult). Stimulus duration is 0.5 s, all other task epochs have variable duration. b. Virtual head fixation algorithm, condition is active for portion of trial marked by green line in A. c. Proportion of trials completed increased as animals were trained on head fixation. Across animals trained on head fixation after learning the visual rule, the mean proportion of completed trials on day 1 of training was 0.498; this increased to 0.679 by day 9. Black trajectories denote animals whose neural recordings were included in this dataset, gray trajectories denote animals who were trained but no recordings were performed. d. Animals typically reached stable performance above 90% accuracy on easy trials in fewer than 30 sessions (median = 16 sessions, +/-10 std). Color scheme as in c. e. Psychometric performance on single sessions after reaching performance criterion on easy stimuli, prior to neural recordings, for each animal included in neural dataset. Error bars indicate standard error of the mean.

The spatially distributed stimuli were designed to exploit the retinotopic organization in V1, but neural responses would only be interpretable if the stimulus could be oriented in a reproducible manner with respect to the animal’s visual field over trials. We therefore additionally required animals to fulfill a head position criterion prior to and throughout the duration of stimulus delivery. We did not control for eye position because we reasoned that the small amplitude eye movements made by rats, which are reduced further when the head is stationary (Wallace et al. 2013), would not impact the low spatial resolution (upper versus lower) at which animals were required to discriminate. Instead of a physical head fixation protocol (Scott et al. 2013), we developed a non-invasive software-based method to virtually constrain the viable head positions at the stimulus viewing port (Figure 1b). We used Bonsai open source software to continuously acquire and segment online video of the behavior chamber (Lopes et al. 2015). Upon trial initiation by the animal, we measured the size and relative position of the animal’s ears in predefined regions of interest (adjusted on a per-animal basis). As long as both size and distance criteria (in both x and y dimensions) were met, the trial was allowed to continue. If any criterion was violated prior to or during stimulus presentation, the trial was aborted and a short time out was delivered. We trained animals to fulfill this postural criterion immediately following acquisition of the decision rule. Rats learned to adjust their head position over the first few sessions of head position training, improving their proportion of successfully completed trials (Figure 1c).

We trained 17 rats to perform this discrimination task, reaching a level of 90%+ accuracy on easiest trials over the course of 16 (median, IQR=16.75) sessions. Of these, 12 animals were trained to maintain head position, and recordings in V1 were made from 5 of these animals. Choice accuracy varied with stimulus difficulty, producing psychometric behavior within and across sessions (Figure 1d).

### Diversity of responses in primary visual cortex during discrimination behavior

We used 32-channel tetrode drives to record putative single unit activity in V1 during this visually-guided (Figure 2a) decision task in order to understand the extent and specificity of task-related information available to this early stage in the visual pathway. We recorded neuronal responses in V1 from 516 units in 5 rats. In what follows we analyze responses from well-isolated single units (n=407), defined as those with consistent, large-amplitude waveforms and fewer than 1% ISI violations. The peak mean activity of an individual unit could occur at any point during the trial, with an enrichment of units showing maximum activity during the movement epoch (Figure 2b,d). The activity patterns were similar in multiunit activity (n=109, Supplementary Figure 3).

**Figure 2.**
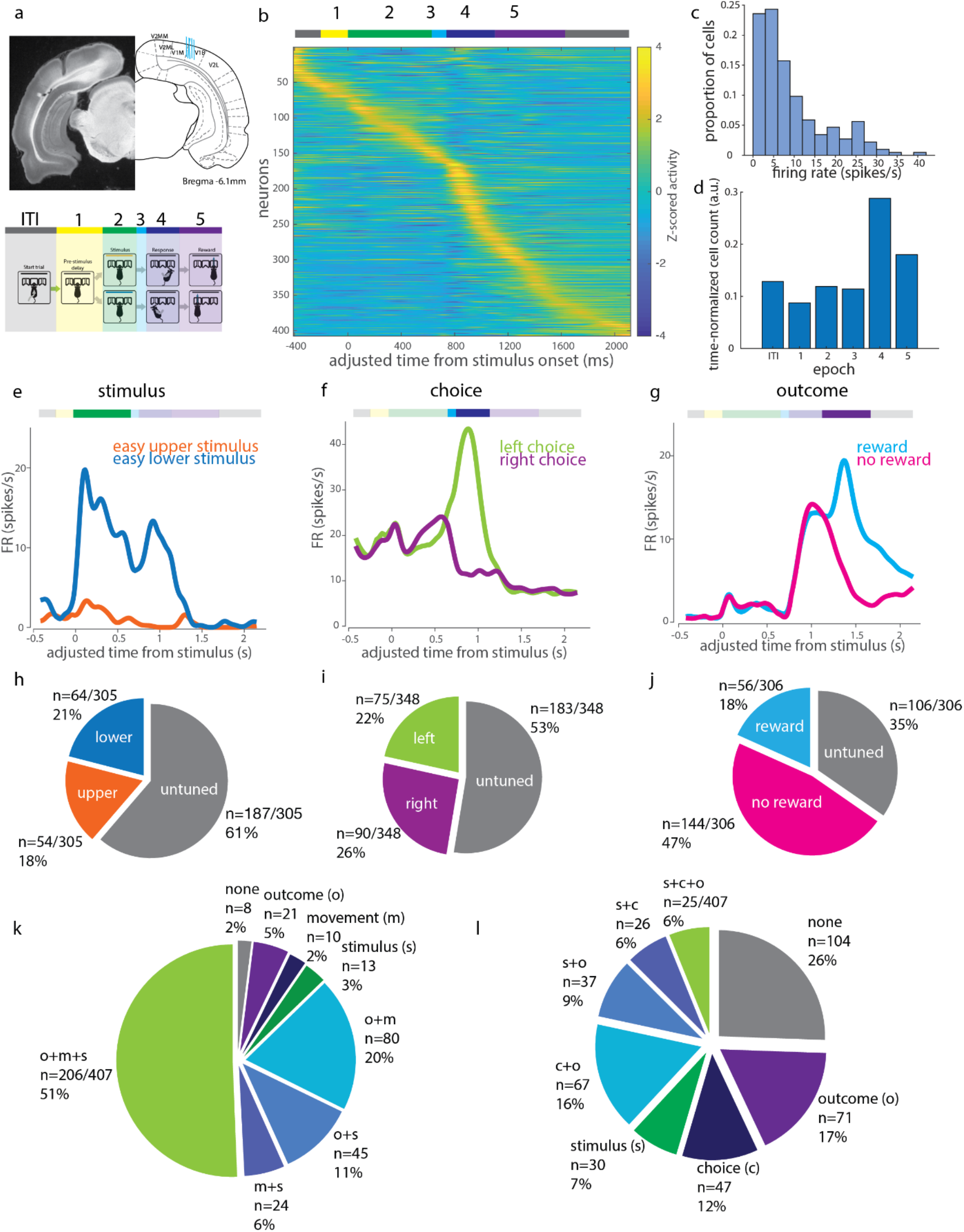
Tuned representations of several task features during visual discrimination by V1 single neurons. a. Recording sites, and definitions of task epochs used in analysis. Schematic adapted from Paxinos and Watson (2007). V1M: primary visual cortex (monocular); V1B: primary visual cortex (binocular); V2L: secondary visual cortex, lateral area; V2MM: secondary visual cortex, mediomedial area; V2ML: secondary visual cortex, mediolateral area; ITI: intertrial interval. b. Mean trial-aligned z-scored activity for all single units in the cloud of dots task (N=5, n=407) spans the duration of the trial. Adjusted time aligns all trials to the same time axis to allow pooling of variable length epochs (see Methods). Task epochs as denoted by colored bar above. c. Firing rate distribution of putative single units. d. Proportion of single units displaying peak activity in each epoch, normalized to the mean duration of each epoch. e. Example neuron preferring stimuli with more dots in the lower half (“lower preferring”). f. Example cell displaying left choice preference during movement period activity. g. Example cell displaying reward preference during outcome epoch. h. Proportion of visual location tuned cells in recording dataset. i. Proportion of choice direction tuned cells. j. Proportion of reward tuned cells. k. Proportions of cells with significant modulation of activity (paired t-test of epoch rates within trials, p<0.05) during stimulus (s, green), movement (m, blue), or outcome (o, purple) epochs compared to pre-stimulus baseline (epoch 1 from panel b). l. Proportion of all single units (n=407) tuned to some combination of stimulus (s), choice (c), and outcome (o) across epochs.

We first quantified the tuning properties of single units to sensory and non-sensory task features during different task epochs. For each epoch of interest, we limited our analysis to single units firing more than 1 spike/s on average during that behavioral epoch. As a result, the set of single units included for each epoch differed slightly (for example, a neuron that fired during stimulus presentation but was silent during movement would be included in stimulus epoch tuning analyses but not movement epoch tuning analyses; see Methods for details). For each feature of interest (stimulus identity, choice side, outcome), we defined a selectivity index (si) to compare the activity evoked by different conditions within a given task epoch:

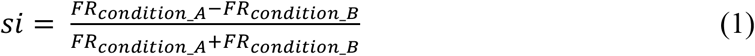

where conditions A and B refer to the two conditions being compared. In the case of stimulus selectivity, for example, F_Reasy lower stimulus_ refers to the firing rate for the 0.5s following stimulus onset when an easy lower stimulus was presented. Comparing the observed selectivity indices to the distribution of indices calculated from the shuffled label control, we identified 39% (118/305) of the single units that were active during the stimulus epoch as significantly stimulus selective (Figure 2h).

We also observed many neurons with above-baseline activity during task epochs other than the stimulus epoch (Figure 2b, k). Activity in later task epochs was often tuned to non-visual task variables such as choice side and outcome. For example, we observed units that preferentially fired during the movement epoch to one choice side over the other, and units whose activity during the outcome period was modulated by reward delivery (Figure 2f,g). Applying the selectivity index analysis to the choice epoch, we found that 47% (165/348) of single units that fired >1 spike/s in this epoch had choice side selectivity across all difficult trials, and thus had “robust choice selectivity” (Figure 2i), while 72% (250/348) were significantly side selective compared to shuffled data controls on at least one trial condition. During the outcome epoch, 66% (200/306) had reward outcome selectivity (Figure 2j). Choice tuning was also significant in a sizeable proportion of units during the outcome epoch (42%, 127/306). Many neurons were selective for combinations of these three features across epochs (Figure 2l). Thus, choice and outcome strongly modulated single neuron activity in V1 during later task epochs, during which many neurons had their peak activity.

We then asked how the specificity of the stimulus-evoked neuronal responses compared to the animals’ behavior. Across the population, the firing rates during the stimulus period were typically modest (mean 7.2 +/− 7.8 spikes/s, median 4.7 spikes/s, Figure 3a), and only a minority (39%) of neurons that were active (>1 spike/s) during the stimulus presentation were selective for upper vs lower stimuli. Of those that were selective, most were weakly selective: Only about 1% of neurons (4/305) had a selectivity index greater than +/− 0.7 (Figure 3b). Across the population, no single unit matched the sensitivity of the animal’s performance on the corresponding session (Figure 3c). We also assessed the trial-to-trial variability in stimulus epoch firing predicting the animal’s choice, by using either a selectivity index or ROC analysis to estimate choice probability. Consistent with previous reports in primates (Nienborg and Cumming 2006), choice probabilities, calculated as the selectivity for future choice from stimulus period activity for a given stimulus condition, were low in V1, with only 2% (5/305) of cells having significant choice probabilities during presentations of difficult stimuli, relative to a shuffle control (Figure 3d). Choice probability calculated using ROC analysis produced similar results (7/305, Supplementary Figure 3a). Thus, activity during the stimulus period reflected the true stimulus more than the perceived stimulus or upcoming choice.

**Figure 3.**
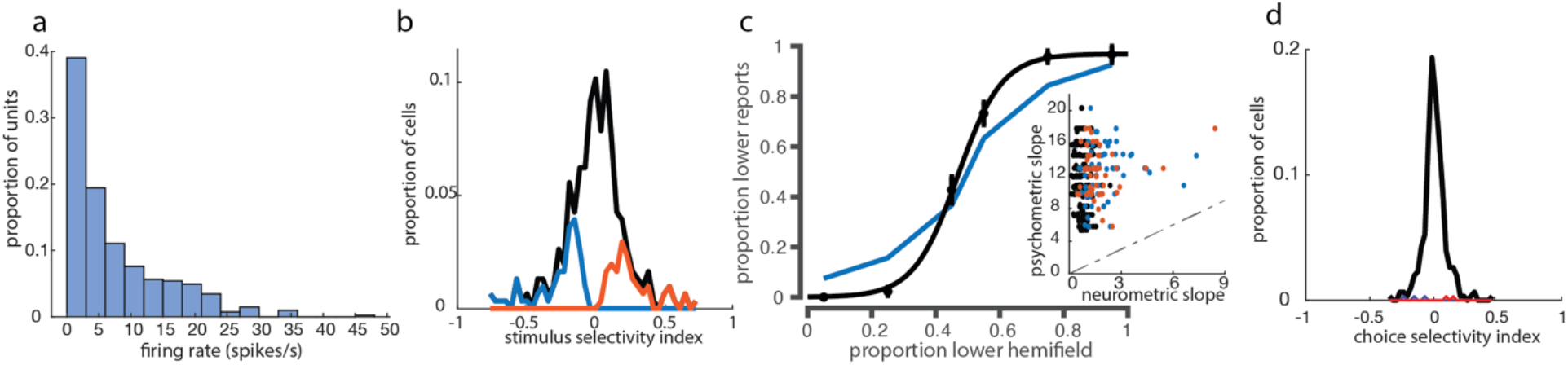
Stimulus epoch activity elicited by “cloud of dots” stimulus is spatially tuned, but less accurate than the animal’s behavior. a. Firing rate distribution across putative single neurons during stimulus epoch. b. Distribution of stimulus selectivity index across all cells active in the stimulus epoch. Blue (lower-preferring, 64/305) and orange (upper-preferring, 54/305) histograms denote cells with significant stimulus selectivity, compared to a shuffle control. c. Comparison of psychometric (black) with neurometric (blue) curve for best lower-preferring cell. Inset: Comparison of psychometric and neurometric slopes across all single units used for stimulus selectivity analysis. Dashed line indicates unity line. d. Selectivity index-based choice probabilities in V1 single neurons (see Methods). Cells with significant choice probabilities are shown in blue (3/305) and orange (2/305).

To further understand non-sensory drivers of activity in V1, we asked whether non-sensory tuning was purely transient, arising only at the moment of the non-sensory event, or whether non-sensory task parameters could exert a persistent influence that spanned trials. We found that some cells were modulated by previous trial parameters, such as whether the previous trial was rewarded, and which choice port was selected in the previous trial (Figure 4a). Such response profiles indicated that choice and outcome tuning do not only influence V1 activity transiently and instantaneously, but rather can be represented in a sustained or history-dependent manner within single cells.

**Figure 4.**
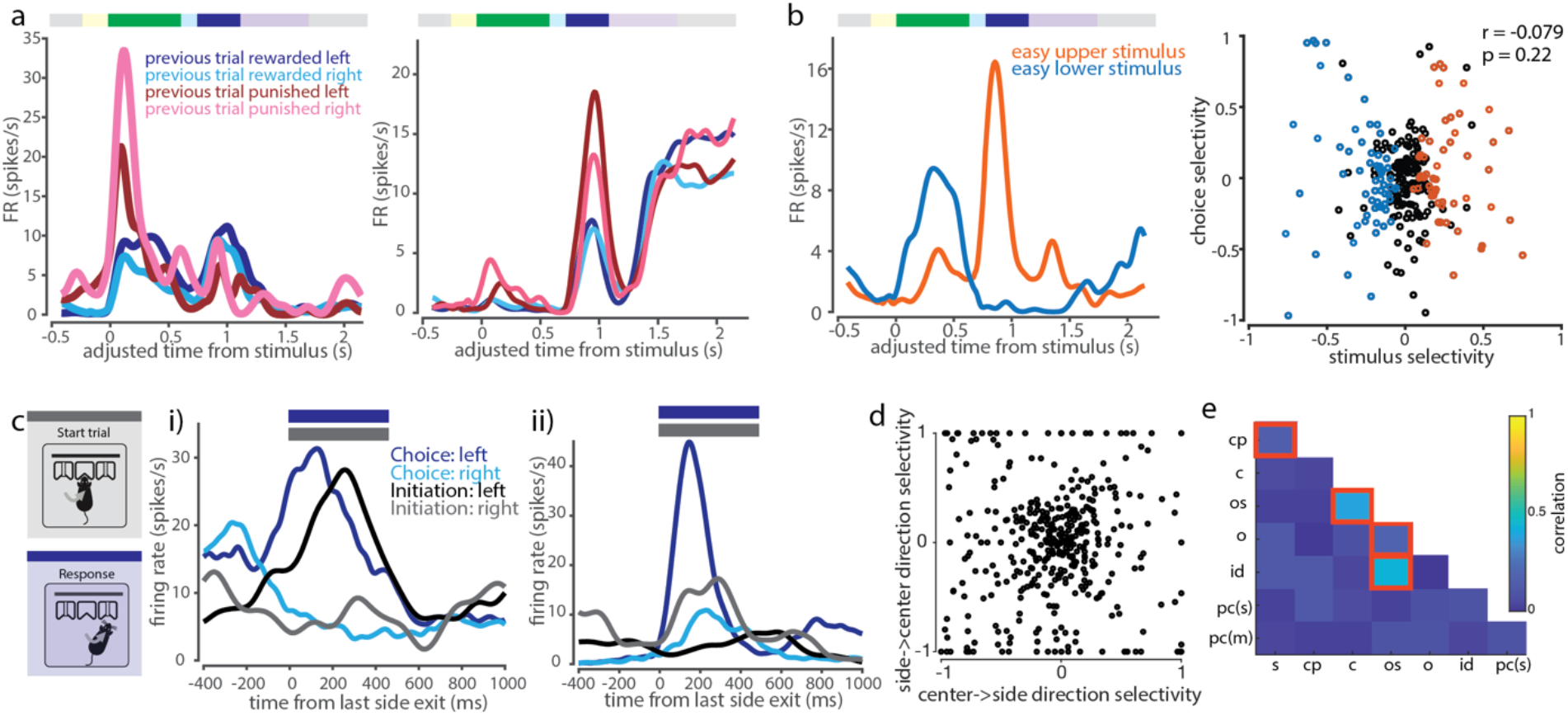
V1 single neuron tuning to non-sensory task variables. a. Example cells showing modulation of task-related activity by previous trial behavioral variables during stimulus and/or choice epochs. b. Left: Example cell showing anti-coherent tuning between stimulus and choice epoch. Right: No significant correlation between stimulus and choice selectivity across cells. c. Comparison of between-port movement responses within movement-responsive cells (initiation epoch, grey, versus choice epoch, blue). i) Example cell with similar leftward-preference during both task epochs. ii) Example cell with varying side preference and amplitude of movement side-selective responses between initiation and choice epochs. d. Side-selectivity index of between-port movements is uncorrelated between choice and initiation movements. e. Selectivity indices across pairs of task features are mostly uncorrelated within neurons. Highlighted squares indicate pairs of features that are significantly correlated (p<0.05, Bonferroni corrected for multiple comparisons). Legend: s = stimulus, cp = choice probability, c = choice, os = outcome side, o = outcome, id = initiation direction, pc(s) = previous choice (stimulus period), pc(m) = previous choice (movement period)

We then asked if there is a systematic relationship between stimulus preference and choice preference in single units. About a fifth (21%, 51/239) of units tuned to either stimulus or choice were tuned to both. However, co-tuning could not be predicted from task contingencies, with tuning opposite to the reinforced association in about half of these neurons (47%, 24/51; Figure 4b). Across the population, we found no correlation between stimulus and choice side selectivity indices (Pearson correlation, p=0.22). Thus, single neurons encoded combinations of stimulus and choice, including combinations that differed from task contingencies reinforced during training.

Similarly, we compared the movement responses elicited during the two between-port movements in our task: the center-to-side choice movement, versus the side-to-center trial initiation movement. We found both cells that displayed similar tuning preferences and response dynamics across the two movements and cells that had different response amplitudes or tuning preferences (Figure 4c). For this analysis, we restricted initiation movements to those that were completed in < 0.5s between side port exit and center port entry, corresponding to direct port-to-port movements of similar latency as choice movements. There was no significant correlation across the population between tuning direction and magnitude, when calculated by selectivity index, across these two epochs (Figure 4d). Thus, movement-direction tuning appeared to be modulated by task epoch.

We repeated this correlation analysis for all pairs of task variables using the selectivity measure described above (Eq. 1). There was in general no systematic relationship between tuning preferences: We observed predominantly weak, insignificant correlations between selectivity to most pairwise combinations of task variables, indicating that tuning preferences were largely independent across task features (Fig. 4e).

Taken together, these analyses show that responses in V1 during this task are driven by features not limited to sensory input, but also including movement direction and outcome, sometimes influenced by multiple parameters, such as previous trial features or current task epoch.

### V1 neurons encode diverse, unstructured combinations of stimulus and task variables within and across task epochs

Having observed a variety of single neuron response patterns in V1, we next set out to quantify the relative influence of different task variables on single neuron activity over the course of a trial. To systematically interrogate how task features influenced single neuron activity at different points in the task, we fit a linear encoding model to estimate the relative influence of each task feature on the firing rate *y* of a given neuron during task epoch *i* (Figure 5b),

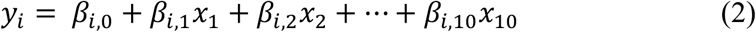

where *i* = 1 … 5 denotes the task epoch; *x*_1_ … *x*_10_ denote the following behavioral variables: stimulus identity (*x*_1_), choice (*x*_2_), reaction time (*x*_3_), movement latency (*x*_4_), choice correctness (*x*_5_), reward delivery (*x*_6_), port last exited on the previous trial (i.e., port visited directly preceding initiation poke, *x*_7_), previous trial choice (i.e., port first visited at previous trial decision time, *x*_8_), previous trial outcome (*x*_9_), and previous trial stimulus identity (*x*_10_); *β*_*i*,1_ … *β*_*i*,10_ are their corresponding weight coefficients within epoch *i*, and *β*_*i*,0_ is the intercept. Note that behavioral variables do not depend on the epoch, as each takes on only one value per trial, i.e. each trial has only one choice side, one reaction time, etc. The model was fit using Lasso regularization with 10-fold cross validation, to derive weights to identify the most informative behavioral variables. We quantified the total variance explained by the model, as well as the relative contribution of each of those variables, by comparing the variance explained by the model when including versus excluding each variable.

In previous analyses above (Fig. 2) we observed that a larger fraction of single neurons in V1 responded during choice and outcome epochs than during the stimulus presentation. Consistent with this, we found that the model also explained a larger total proportion of the variance of choice and outcome epoch activity (mean variance explained of 0.19 and 0.25, respectively, Figure 5b), compared to the stimulus epoch (mean variance explained of 0.09; distributions are significantly different by the Kolmogorov-Smirnov test, p<10^-14^ for both). Furthermore, within the stimulus epoch, we found more total cells whose activity was better explained by one of several previous task features, such as previous choice, outcome, and exit port side, than by current stimulus identity. Thus, single neuron firing variability was consistently better explained by non-stimulus task variables, over the course of the trial and even during stimulus presentation.

**Figure 5.**
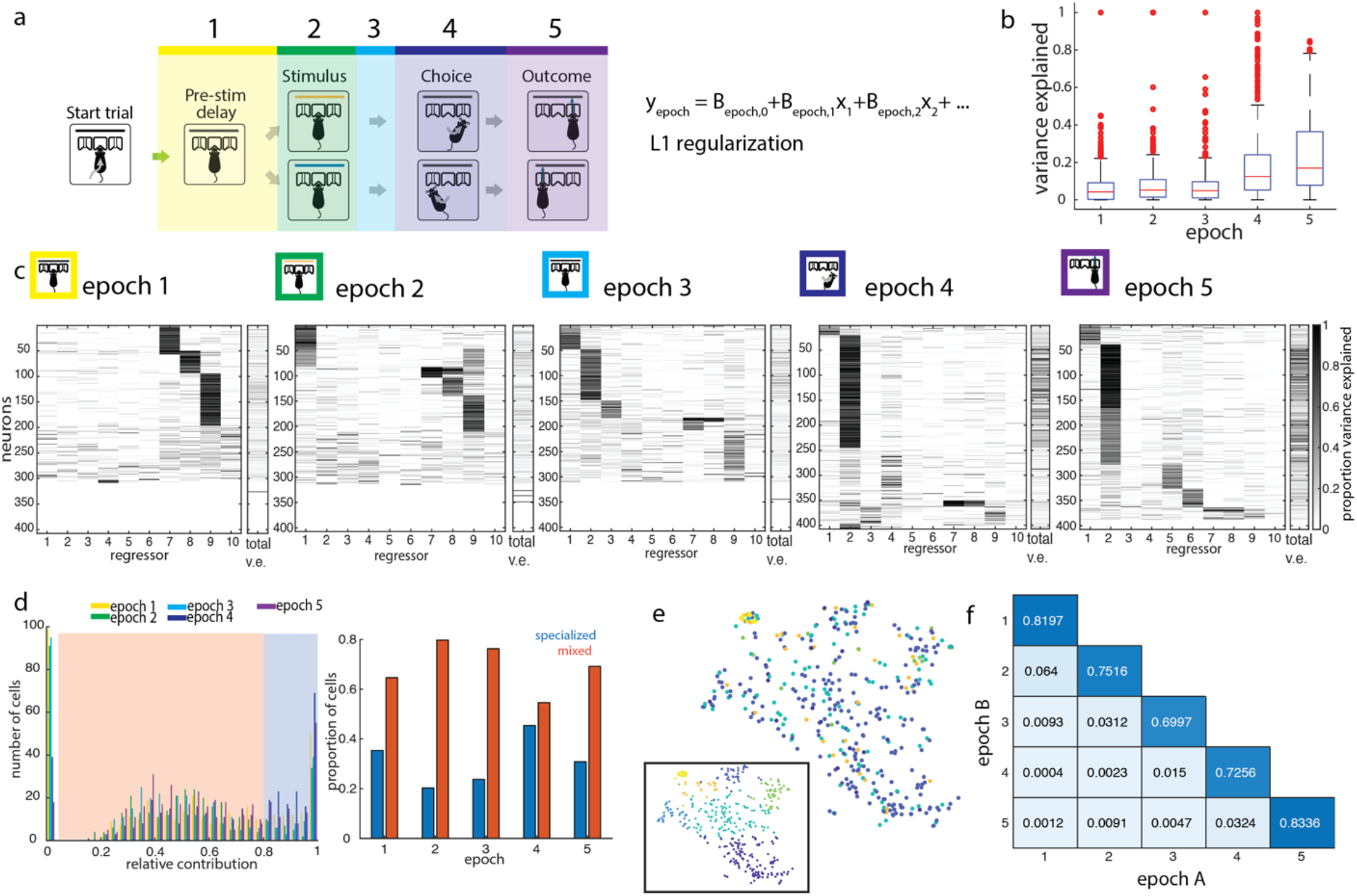
Single neurons represent combinations of task features within and across task epochs. a. Design of linear encoding model. Trial divided into 5 epochs, as marked. Linear model was fit using 10 task parameters to predict trial-by-trial firing rates within epochs: 1) stimulus, 2) choice, 3) reaction time, 4) movement latency, 5) correctness, 6) reward delivery, 7) previous trial last port visited, 8) previous trial choice, 9) previous trial outcome, 10) previous trial stimulus. b. Box and whisker plot of total variance explained by the model, by epoch. c. Relative variance explained by individual regressors in the linear encoding model, by epoch. Total variance explained for each neuron is shown in the rightmost column in each epoch. The left 10 columns show the proportion of the explainable variance attributed to each regressor for each neuron (darker shading = higher proportion of total variance explained, see Methods). Neurons (rows) are clustered and sorted within epochs. In some units, single regressors dominate the explainable variance, while in others, multiple regressors contribute to the encoding model, revealing the presence of both “specialized” and “mixed” encoding by cells during each epoch. d. Distribution of maximum contribution by a single task parameter to predictions. Thresholding at a relative contribution of 0.8 separates cells into “mixed” (orange shading) and “specialized” (blue shading) encoding profiles. Cells with maximum relative contribution near 0 are excluded as not being well-driven by any of the regressors. Right: Proportions of specialized versus mixed encoding cells across epochs. e. t-sne embedding of encoding profiles of single units in the outcome epoch, clustered by cluster identities from the choice epoch. Inset shows the same embedding, clustered by outcome epoch cluster identities. Color denotes cluster identity. f. Cluster goodness-of-fit measure (adjusted Rand Index; see Methods) for all pairwise comparisons of epochs A and B. Clustering different epochs produces fewer shared cluster members than two independent partitions of the same epoch.

Of the activity explainable by our model, we wanted to know whether cells were predominantly “specialized” for encoding a single task variable, or encoded a “mixture” of task variables. Based on the distribution of the most prominent task feature’s contribution to the linear model, we set a cutoff that classified features surpassing a relative contribution of 0.8 as dominating a given neuron’s response, and that neuron was subsequently designated as “specialized” during that epoch. Otherwise, the neuron was designated as having “mixed” representations, with more than one task variable contributing substantially to its activity in that epoch. In most epochs, the majority of single neurons (between 55% and 80%) were driven by a combination of task features, rather than a single feature. The closest ratio was in the choice epoch, where there were almost as many specialized choice-selective neurons as there were neurons encoding a mixture of stimulus, choice, and other movement related features such as reaction time (Figure 5d). Therefore, task information was encoded not by multiple independent groups of specialized cells, but rather by overlapping modulation of the activity of single cells.

The predominantly mixed profiles of neural responses argue against a simple labelled line model, in which each task variable is represented by a particular class of cells receiving input predominantly from a single source. We therefore considered a somewhat more complex model in which neurons within a cell class represent similar sensory and non-sensory variables between them, across epochs, i.e. two neurons that represent the same combination of features in the stimulus epoch will also look similar to one another in their encoding patterns in the choice epoch. To test this, we clustered neurons on the basis of the relative contributions of all task features in a given epoch (e.g. choice epoch), and used these clusters to sort the relative contribution of task features to their activity in each of the other epochs (e.g. outcome epoch, Figure 5e). We found that no distinct clusters emerged in the outcome epoch, when cells were ordered by their cluster identity in the choice epoch. We repeated this for all clustering epoch–test epoch pairs and saw that cluster identity always generalized poorly across all pairs of epochs (Figure 5f). This is reflected in the adjusted Rand Index, a standard measure which quantifies the overlap in cluster membership between two independent partitions, and was much lower for cross-epoch comparisons than within-epoch comparisons. The adjusted Rand Index, which ranges between 0 and 1, is maximized when the same sets of neurons are clustered together in both partitions. Thus, single neurons represent diverse combinations of task variables both within and across epochs, without any evident organization or structure.

### Current and past trial task features can be decoded from V1 population activity

The single neuron encoding patterns we observed suggested that the encoding of task variables was distributed across a heterogeneous V1 population. Such shifting representations at the single cell level may nonetheless underlie stable representations at the population level. We therefore analyzed the information available in populations of simultaneously recorded cells throughout the duration of a trial. First, we used dimensionality reduction methods to inspect the population activity of simultaneously recorded units (both putative single units and multi-unit activity) over the course of single trials (Figure 6a). Activity patterns diverged over the course of the trial on the basis of stimulus identity, choice side, and outcome, and evolved along distinct dimensions during the stimulus, choice, and outcome periods. This suggested that it would be possible to read out these task features from V1 population activity at different points in the trial.

**Figure 6.**
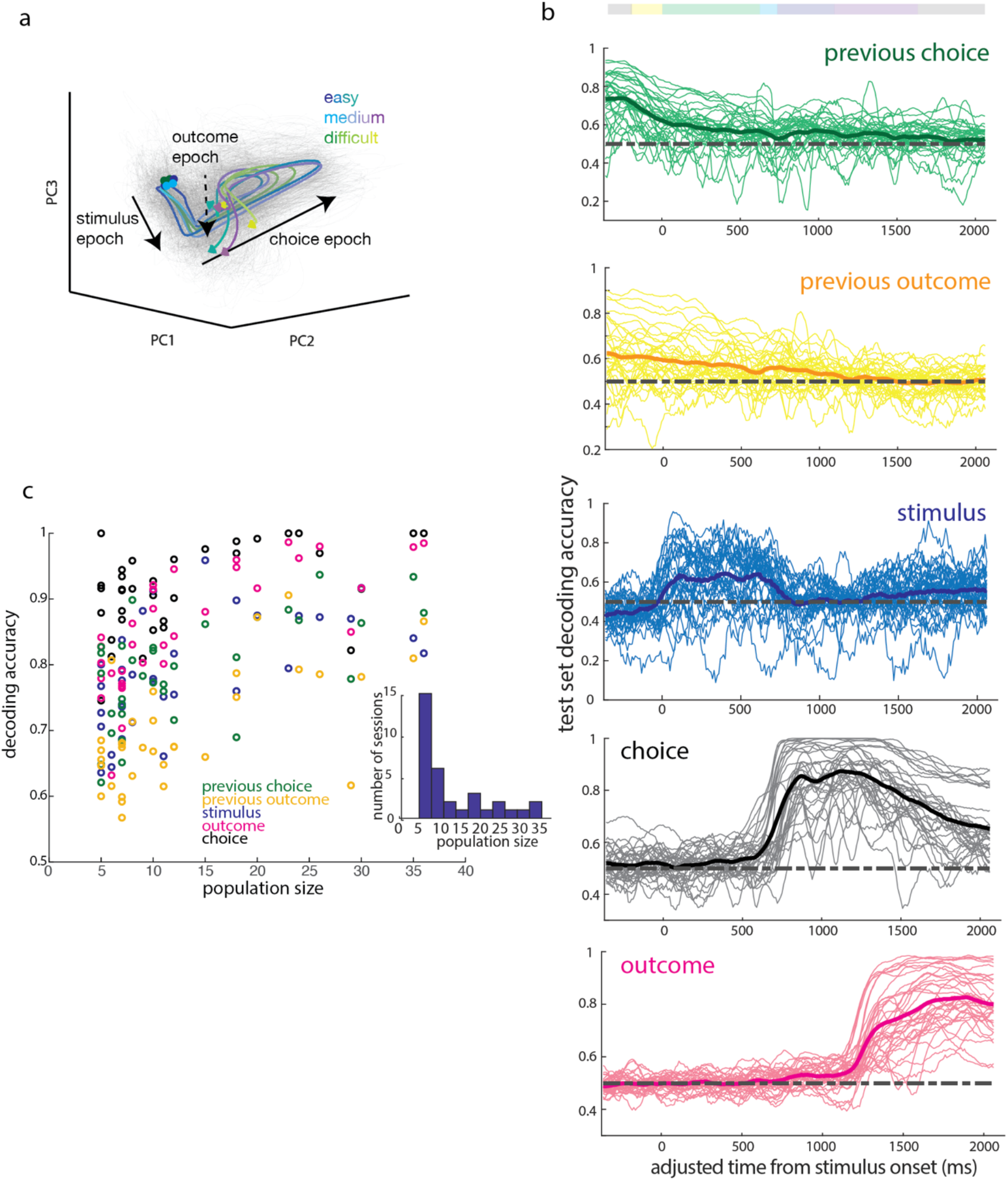
Reliable decoding of task variables across trial duration from trial-by-trial population activity. a. Single trial (grey) and mean (within conditions, colored by trial difficulty) population activity trajectories for an example session, projected onto the first 3 principal components. b. Decoding accuracy in sliding 100ms bins over the course of a trial for features of the previous trial (choice side, outcome), and of the current trial (stimulus category, choice side, and outcome). Decoding accuracy calculated as proportion of test set classified correctly from activity at a given timepoint. Thin lines correspond to individual sessions, while bold lines denote the mean across sessions. c. Maximum decoding accuracy of trial features as a function of population size. Inset: Distribution of population sizes.

To test how well features of the task could be decoded from the population activity at each timepoint, we trained a linear classifier to decode task variables: stimulus category, choice, and outcome, previous choice and previous outcome (Figure 6b). We found characteristic decoding timecourses for each feature. Stimulus category could be decoded primarily during stimulus presentation (see Methods: Decoding (Linear Classifier)). Task features associated with the previous trial, such as previous choice and previous outcome, could be decoded early in the trial, with performance decreasing over the course of the trial. Consistent with this, choice and outcome were readily decoded both during and following their respective epochs. Outcome information could be decoded regardless of whether we pooled missed reward and punishment outcomes, or treated them separately (Supplementary Figure 4a). The timecourse of how well each feature could be decoded from the neural activity was similar across sessions for any given feature, which is reflected in the proportion of sessions with significantly better-than-chance decoding accuracy over the course of the trial (Supplementary Figure 4c-g). Thus, multiple task features could be read out from population activity at each timepoint over the trial, including during early epochs when single neuron activity was less well explained by the previous encoding model.

Decoding accuracy improved on sessions with more simultaneously recorded units, but notably, even the smallest populations included in this analysis (5 units) were able to exceed a decoding accuracy of 60% for most task features (Figure 6c). In addition, classifier performance did not increase substantially with population size beyond about 20 units. Thus, despite the heterogeneity of single neuron activity patterns, task information could readily be decoded by a linear decoder from small V1 populations, with a similar timecourse over sessions.

### V1 representations during visually-independent choice task

The robust task-related representations we observed in V1 could be specific to visually-guided decisions. Alternatively, non-sensory representations might be encoded in visual cortex independently of whether primary visual cortex is required for the decision process. To distinguish these possibilities, we interrogated V1 responses in a new cohort of subjects trained to perform a similarly structured task in which decisions were based on auditory rather than visual stimuli. In this modified task, visual stimuli were presented but not informative for the animal’s choice. Instead, animals were instructed as to the correct choice based on the location of the decision tone, which was presented on the side of the animal corresponding to the correct side port for that trial. The task structure was otherwise identical to that of the visual discrimination task (Figure 7a). During this task, the visual stimuli consisted of randomly dispersed dots over the full extent of the monitor on the majority (70%) of trials. On the remaining trials, animals were presented with one of the two “easy” stimuli from the discrimination task. Animals acquired this task to near-perfection, and their choice profiles were uncorrelated with the distribution of the visual stimulus (Supplementary Figure 5).

**Figure 7.**
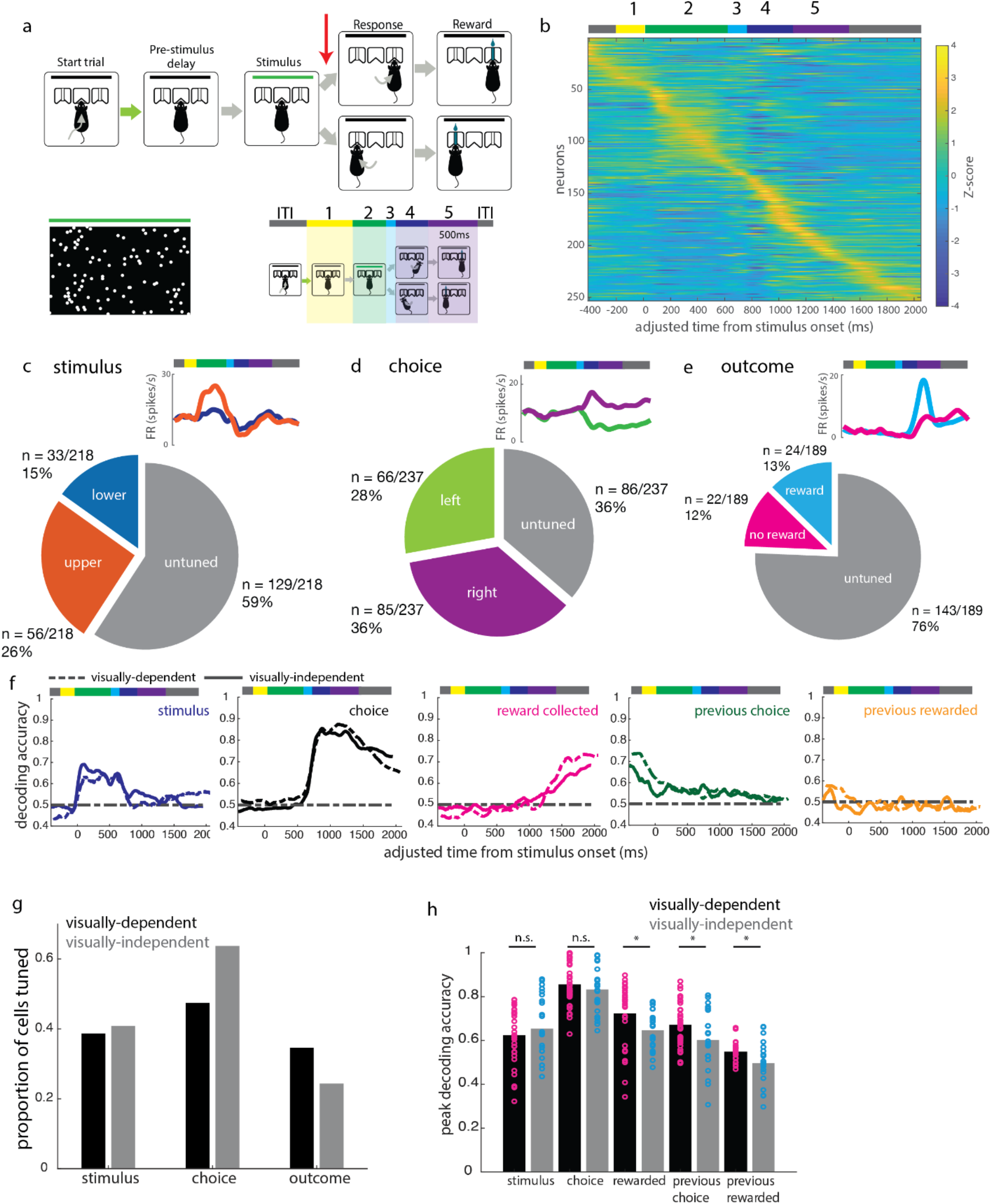
V1 responses during a visually-independent decision task follow similar patterns. a. Task structure is identical to the structure of the visual discrimination task, except that decision tone (red arrow) is presented on one side only. A response to the same side as the decision tone yields a reward. b. Z-scored mean activity of single units, sorted by time of peak activity. c-e. Example neurons and proportion of single units selective for (c) stimulus, (d) choice side, and (e) reward delivery. f. Mean decoding trajectories over visually-independent decision sessions (solid lines) for current trial stimulus, choice, and outcome, previous trial choice, and previous trial outcome. Dashed lines denote mean trajectories during the discrimination task, as shown in Figure 6. g. Comparison of proportion of tuned cells between visually-dependent and visually-independent choice tasks. h. Comparison of decoding accuracy for V1 populations between visually-dependent and visually-independent choice tasks, during the 500ms of the trial with the best performance on decoding of each task feature. Bars (black = visually-dependent task, gray = visually-independent task) indicate mean decoding accuracy across trials, while overlaid points indicate accuracy on single trials. Significant differences in decoding accuracy between the tasks were found for the following non-sensory parameters: outcome, previous choice, and previous outcome (2-sample t-test, * p<0.05).

We recorded from 253 well-isolated single units and 41 multi-units from 2 animals performing this task variant. The trial-averaged activity across the population was similar to that recorded in the visual discrimination task, with the majority of units having their peak firing during or after the movement epoch (Figure 7b). Firing rates were similarly modest, with a mean of 6.5 (+/-4.4 std) spikes/s (Supplementary Figure 6a). Stimulus selectivity profiles were also similar between the two tasks: 41% of single units were stimulus selective in the visually-independent task (Figure 7c,g). The proportion of choice selective cells increased from the proportion of robust choice selective cells in the discrimination task (64% compared to 47%, Figure 7d,g), while the proportion of outcome selective cells decreased. Because errors were rare in this task, we instead compared rewarded versus missed reward trials (i.e. a correct choice where the animal’s choice port nosepoke was too short in duration to trigger a reward). 35% vs 24% of cells in the discrimination and visually-independent tasks were selective between rewarded versus missed reward trials, respectively (Figure 7e,g).

Decoding task features from population activity yielded timecourses similar to those obtained on the visually-guided task, with some differences in peak decoding accuracy. While stimulus and choice decoding were accurate to similar levels as in the discrimination task, the decoding performance for rewarded vs missed reward trials was significantly reduced (Figure 7f,h, Supplemental Figure 7a-c). In addition, the onset of outcome decoding was delayed, compared to in the discrimination task, to after the first 500ms of the outcome epoch. The slower and decreased rise in outcome information is consistent with the execution of different motor programs following reward versus no reward, in the late outcome period. Previous trial choice and outcome decoding accuracy were also reduced during the visually-independent task, further suggesting that the representation of some task features (stimulus, choice) in V1 are robust across task demands, but others (outcome, previous choice, previous outcome) are task-dependent.

Finally, when fitting the same linear encoding model across the two tasks, we found that single neuron activity in the visually-independent decision task was 1) similarly predominantly driven by more than one task feature at a time, and 2) similarly better described at later points in the trial (choice and outcome epochs) than at early points in the trial (including the stimulus epoch, Supplementary Figure 6b-d), as in the visual discrimination task (Figure 5). However, the total proportion of the variance explained by our model was significantly lower in each epoch in the visually-independent decision task, compared to the same epoch in the visual discrimination task (Supplementary Figure 6e), which is consistent with decreased influence of some task features on V1 activity during the visually-independent task.

The comparison of the visually-guided task with the non-visual task reveals that while neural activity in V1 was broadly similar between the two tasks, encoding of the non-sensory task features we investigated here – choice and outcome – were differently affected by the behavioral context. Representations of outcome in single cells and across the population were less prominent in V1 during visually-independent decisions, while representations of choice remained robust. Previous trial features were also less well represented at the population level, further suggesting that processing of non-sensory information in V1 in a freely moving animal depends somewhat – but not entirely – on the behavioral demands related to visual processing.

## Discussion

In this study, we developed a novel visual discrimination task for freely moving rats to study representations in primary visual cortex during freely moving visual decisions. By recording single unit activity during this behavior, we found robust tuning for both sensory and non-sensory task features, and that tuning preferences were distributed and independent of stimulus-choice contingencies. Single cells were more likely to be driven by multiple features in each epoch than a single task feature. Task features could be decoded from small simultaneously recorded populations of units, with previous trial features best decoded early in the trial, and giving way to current trial features as the trial progressed. Finally, many of the tuning patterns described for the visual discrimination task held true during a visually-independent variant of the task, with the notable exceptions of outcome and previous trial task parameters, for which population decoding accuracy was significantly diminished.

To perform these experiments, we developed a virtual head fixation protocol that is noninvasive, compatible with experimental techniques, and learnable without a direct reinforcement signal. This allowed us to restrict the viewing angle of visual stimuli in a freely moving animal, which we combined with well-defined choice reports and measures of behavioral timing. This system allowed us to impose a real-time postural criterion into training protocols for our task. At the time these experiments were initiated, deep-learning based pose estimation algorithms were not yet available for implementation of real-time video tracking and reactive control of behavioral hardware (Mathis et al. 2018), although they have since been developed (Forys et al. 2020, Kane et al. 2020) and could be used to refine this training approach.

The presence and organization of task representations in visual cortex have implications for the computations that can occur locally and in circuits involving V1. In frontal and parietal cortices, where representations of diverse task-related variables are more frequently studied, there is debate as to whether representations are randomly assorted across neurons, or organized into discrete classes, with potential implications for downstream decoding (Rigotti et al. 2013). Recent work has identified distributed encoding profiles in both cortical (Levy et al. 2020) and subcortical brain regions. In VTA dopaminergic neurons, different degrees of specialization arise in different task epochs (Engelhard et al. 2019), and the specific variables encoded by a given neuron also varies across task epochs. Here, we observed similar complexity in the encoding patterns in a primary sensory cortical area, V1, with cells tuned to the same variable during one task epoch later representing different variables between them in a later epoch, with uncorrelated tuning preferences. Within individual epochs, representations of a given task feature were distributed across the population. In the stimulus epoch, single neurons were less accurate than the animal at classifying the incoming stimuli, and over the trial, both sensory and non-sensory task parameters were decoded better with increasing neural population size of up to ~20 units. Taken together, our results suggest that the primary visual cortex may share some organizational principles with frontal and parietal areas, in that task feature representations are distributed across neurons.

One striking observation was that the ability to decode task features from V1 populations could extend well past the event’s duration, into the next trial, during visually-guided but less so in visually-independent decisions. This argues against the possibility that non-sensory responses in V1 merely reflect an instantaneous “echo” of a brief event such as a motor command. Rather, visual cortex has the ability to carry sustained representations of different task parameters, in a task-dependent manner. Recent work has suggested that non-sensory responses in V1 help shape sensory processing by influencing the correlation structure and population activity space (Osako et al. 2021). Here, we found that sensory processing demands influence which non-sensory correlates are available in V1.

Which characteristics of the task influence whether V1 will carry these non-sensory representations? Because our two tasks are identical in trial structure, but differ in whether the animal is required to use a visual stimulus to guide its behavior, we suspect that the relevant characteristic is whether the task requires visual processing. Another possibility is that V1 task representations depend on the overall difficulty level of the task, i.e. whether difficult (perceptually ambiguous) trials are included. Either way, flexible routing of task-related information through V1 suggests that non-sensory representations may serve a task-dependent computational role. For example, previous-trial parameters may support learning of expectations about the structure of the task and stimulus space.

The stimulus-choice associations that animals were trained on were not reflected in the cotuning preferences of single cells (Fig 4). This was surprising in light of previous studies (Poort et al. 2015, Puscian et al. 2020), in which coherence between visual encoding and behavioral response emerged over training. There are a number of differences in these tasks that could account for these differences. First, in previous studies the visual stimulus and the appropriate response overlapped in time, whereas in our task they were temporally separated. Second, in previous studies the stimuli and eventual outcome were deterministically paired (e.g. only one stimulus could lead to reward), whereas in our task both stimulus categories were equally likely to lead to reward. Finally, there are differences in the V1 neuronal populations sampled: the previous work used two photon imaging, which predominantly samples neurons in layer 2/3, whereas in our study we used tetrodes and thus sampled deep layers as well. Layer 5 neurons in V1 tend to have larger and more complex-like receptive fields (e.g. wider orientation tuning curves, (Niell and Stryker 2008)), and it has been hypothesized that layer 5 V1 neurons may carry out distinct computational functions compared to neurons in layer 2/3 (Keller and Mrsic-Flogel 2018). Future work delineating the behavioral limits where coherence between sensory and non-sensory representations no longer develops may provide clues to how visual cortex processes non-sensory information to support different tasks.

In the context of recent work, our study adds to the growing evidence that the range of responses measured in visual cortex extends far beyond visual stimulus-driven activity. In particular, we contribute evidence for diverse, distributed task representations in V1 in freely moving rodents, complementing the growing literature on V1 activity in awake head-fixed rodents.

## Materials and Methods

### Animals and surgical procedures

All procedures were conducted in accordance with the institutional animal use and care policies of CSHL and NIH. 8-10 week old male Long Evans rats were obtained from Taconic Biosciences and Charles River, and started training after reaching at least 10 weeks of age. Rats were pair-housed until implantation of the microdrive, after which they were singly housed, in a reverse 12h light/dark cycle. Implant surgeries were performed under 2% isoflurane anesthesia. Custom-built microdrives were implanted according to stereotaxic coordinates, with the tetrode bundle targeted to left binocular primary visual cortex (bregma – 6.1mm AP, +4.5 mm ML).

### Task design and behavioral system

Custom behavioral chambers consisted of three ports attached to a clear wall panel through which a monitor was visible to the interior of the behavioral box. Interruption of an infrared beam inside the ports were used to determine timing of port entry and exit. We used the Bpod system (Sanworks, NY) to implement the behavioral state machine. The task structure was as follows: animal entry into the center port triggered the beginning of a pre-stimulus delay. The variable pre-stimulus delay was drawn from an exponential function with a mean of 0.3 s. Following this delay, a 500ms fixed time stimulus was delivered through Psychtoolbox (Brainard, 1997; Pelli, 1997; Kleiner et al, 2007). A 200ms fixed post-stimulus delay separated the stimulus off trigger from the decision tone. Any withdrawal from the center nosepoke at any point between the pre-stimulus delay initiation and the decision tone delivery led to a missed trial and a 2s time out. After implementation of the head position protocol, a missed trial could also be triggered by a head movement while in the center port during this peristimulus period. After the decision tone, the animal was given 3s to make a decision by poking into a side port. A 20 μL reward was delivered following a 50ms nosepoke into the correct port. A correct choice report that did not fulfill this duration requirement did not trigger reward, but no punishment was delivered either. No intertrial interval was specified following correct (either rewarded or missed reward) trials. A 1s punishment tone (white noise stimulus) and a 5-6s time out followed an incorrect choice.

The Psychtoolbox toolbox was used to generate and deliver visual stimuli and auditory decision and punishment tones. For each stimulus, 30 frames were delivered at 60Hz refresh rate, with stimuli randomly distributed across each frame according to the stimulus condition on that given trial. For the discrimination task, the stimulus consisted of two subregions of equal size, separated by a thin boundary region where no dots were ever present. For the stimulus-independent task, dots were presented across the full extent of the display. Across all frames, dots were presented at 1% of all possible locations. For the discrimination task, the less dense subregion on each frame was given the number of dots drawn from a Poisson distribution centered on the lesser mean dot value of that stimulus condition. The denser subregion was given the complementary number of points. Therefore, every frame had the same total number of individual dots. Each dot location contained a round white dot that subtended about 3° in visual space. For the stimulus-independent task, the stimulus period was increased to 700ms, so 42 frames were delivered on each trial. A luminance detector module (Frame2TTL, Sanworks) reported luminance changes during each trial and the onset of stimulus delivery by detecting a reporter pixel which flickered on/off with each frame update.

### Head position control

We implemented the closed-loop head position condition using Bonsai, a reactive programming software (Lopes et al. 2015). Bonsai was given video input from a webcam (Logitech) mounted above the animal at a 70° angle. This video input was binarized and regions of interest (ROIs) were defined on a per-animal basis from this field of view. These ROIs were centered on the position of each ear, such that the ear would entirely fall within the ROI when properly aligned. Built-in Bonsai functions carried out contour mapping of the image within each ROI, and filtered viable objects on the basis of size. The centroid positions of the resulting objects were calculated, and if their distance did not exceed a threshold of 10-15 pixels, a binary signal representing the animal’s successful alignment was sent to the behavioral state machine. This condition was only tested for when the animal was in the port to prevent spurious detections or noise caused by background (e.g. behavior rig floor) objects. The algorithm performed a moment-to-moment “and” computation on the comparison between the x values, the comparison between the y values, and the input trigger to output a binary trigger back to Bpod. The continuation of the Bpod states depended on the continuous on-state of this trigger. To ward against fast software- or camera-generated errors from producing false negatives, a short 50ms grace period followed every on-off transition of the trigger. If during this grace period the trigger returned to the on state, the trial was allowed to continue; otherwise, it was aborted.

### Extracellular recordings

Tetrode drives were custom-built using Omnetics 36-channel EIBs and custom 3D printed drive skeletons. Each drive contained 8 tetrodes and 1 reference tetrode that travelled together in a single bundle. Subjects were implanted with tetrode drives under 2% isoflurane anesthesia following successful acquisition of both the visual discrimination (where applicable) and the head position requirement.

We used the Intan-based OpenEphys recording system to acquire neural signals. Four of the seven animals reported here required light anesthesia to facilitate attachment of the recording tether (2/5 on the discrimination task and 2/2 on the visually-independent choice task). These animals were given 15 minutes to fully recover before the task began. After each recording session, tetrodes were lowered by 40-80μm. Recordings were made until tetrodes reached a depth of 1.5mm. We electrolytically lesioned at the tetrode tips, after which animals were sacrificed and brains were recovered for histology.

Spike times were extracted through semi-automated spike sorting using Kilosort software on the raw continuous recording traces. The data was bandpass filtered and the mean across all channels was subtracted from all traces to remove any common noise events. We performed manual curation of detected spikes on the basis of their: amplitudes, waveforms, auto- and crosscorrelograms, firing dynamics over the session, and clustering in feature space. We further restricted single cell representation analyses to units with refractory period (2ms) violations of less than 1%. All analyses were performed in Matlab.

### Time adjustment / neural data preprocessing

Individual trials varied slightly in duration due to variable durations of pre-stimulus delays, reaction times, and lengths of stay in reward ports. For all analyses that did not rely on mean epoch firing rates, to allow comparisons of firing rate trajectories over trials and sessions, e.g. in figures 2, 6, and 7, we first “stretched” individual trials to a common timecourse across all recorded sessions. We sampled individual activity traces at regularly spaced timepoints within each epoch, then mapped those sampled points back to the mean trial timecourse.

### Selectivity analyses

To find the selectivity of a cell’s firing during various task epochs, a selectivity index was calculated on the mean firing rates between pairs of trial types defined by the task parameter of interest. We defined selective cells as those whose selectivity index exceeds the 95% bounds of a shuffle control distribution. The shuffle control distribution for a given cell was built by calculating the selectivity index across 1000 shuffles where the trial labels (e.g. upper or lower stimulus) were shuffled relative to the single trial firing rates for that cell. We carried out the same analysis to define movement side-selective cells during the choice epoch, and reward-selective cells during the outcome epoch. For each epoch of interest, of the total single units (n=407), only those with an average firing rate of more than 1 spike/s during that epoch were included in this analysis (stimulus epoch: 305 cells; choice epoch: 348 cells; outcome epoch: 306 cells).

Selectivity analyses in figures 2–4 were calculated for variables including: stimulus (more upper dots vs more lower dots); choice (left port entry at decision tone vs right port entry); choice probability (eventual choice, neural activity during stimulus delivery); outcome (rewarded vs not rewarded); outcome side (left port during outcome epoch vs right port); initiation direction (approach to center port from left vs right port); and previous choice (left vs right port selected on previous trial).

### Neurometrics

ROC analysis was performed using the Matlab *perfcurve* function, using task variable as a binary label, and mean single trial firing rates in a given task epoch as the scores. To build the neurometric curve, we applied ROC analysis at each of the 3 stimulus difficulty levels presented, and took the area under the curve as the cell’s ability to discriminate between the two easy, the two medium, and the two difficult stimuli. These values were mirrored across the 50% point of the decision axis to estimate the full psychometric curve. For comparison of the slopes of the neurometric and associated psychometric curves, we fit a logistic function to the 6 points from the auROC analysis, and a second logistic function to infer the psychometric function from the choice behavior, and compared the slope parameter from these two fits.

### Linear Encoding Model

We trained a linear model to predict the firing rate during each epoch given the set of behavioral predictors. Binary variables (e.g. choice, correctness, and reward delivery) were coded as values of −1 and 1. Continuous-valued variables (e.g. reaction time and movement duration) were z-scored over the session. Stimulus identity took on a value between −1 and 1 which represented the comparison strength in the stimulus (proportion of dots_lower_ – proportion of dots_upper_). We used Lasso regularization, setting lambda to minimize the deviance across validation sets. We carried out this model optimization using the Matlab *lassoglm* function, with 10x crossvalidation. Variance explained by the model predictions (*η*^2^_*model*_) was used as a measure of model fit, calculated as:

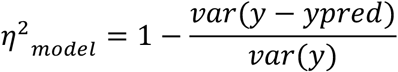

where *y* is the measured firing rate, and *ypred*is the firing rate predicted by the model. Proportion of variance explained for predictor *i* was used as a measure of the predictor’s contribution to the model, calculated as:

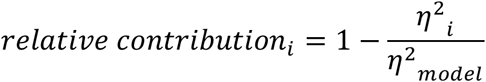

where *η*^2^_*i*_ is the variance explained by the model lacking the predictor *i* (i.e. the weights for predictor *i* are set to zero after training), and *η*^2^_model_ is the variance explained by the full model.

Neurons were clustered by their encoding weights using k-means clustering with the number of clusters k determined by maximizing the adjusted Rand Index (ARI), a measure of clustering stability, as a function of number of clusters. We first removed all zero vectors (corresponding to cells that were not explained by the task variables), then computed ARI as the average similarity of 500 pairwise comparisons of independent clusterings of the encoding weights in a given epoch, for k = 2 to 10 clusters. In order to compare stability of clusters across epochs, we chose to use a constant number of clusters across epochs, so we pooled the ARI across epochs to find the peak of the mean curve as a function of k. This gave an optimal k of 6 for clustering cells with non-zero weight vectors, then for the sake of comparison between epochs, we added back the final “cluster” of zero weight vector cells for that epoch to make a total of 7 clusters per epoch.

Comparison of clustering similarity across epochs was measured using the ARI as a measure of pairwise similarity of the clustering between pairs of epochs. This similarity was computed including the cells with zero weight vectors.

### Decoding (Linear Classifier)

Population activity at a given timepoint was expressed as a vector of mean rates over a 100-ms bin centered at the timepoint of interest, for all units recorded on a given session. To estimate the timecourse of activity, activity in 100-ms sliding bins were calculated every 10ms. To visualize activity trajectories over the trial, principal components decomposition was applied to the population activity matrix, and the activity was projected onto the first 3 principle components.

To assess the amount of information available about a given task variable in the population activity for downstream readout, we trained a linear classifier using the Matlab function *fitclinear* with 5-fold cross validation and lasso regularization on the activity patterns and task variable labels from 90% of valid trials (more below), and assessed the accuracy of predictions on the held out 10% of trials. We repeated this modelling 100 times to assess stability of the trained models. We trained the classifier independently at each timepoint, and then compared the learned weights across timepoints and across models. The weights were highly consistent across trained models at a given timepoint, but varied for a given neuron over the course of a trial.

Valid trials were defined as trials on which subjects completed the full trial (through stimulus presentation and the post-stimulus delay). To assess choice decoding, we further restricted the trials used to difficult trials, where stimulus discriminability was low and choice profiles approached chance. To assess stimulus decoding, we used trials where the easiest stimuli were presented, to facilitate a one-to-one comparison between the two tasks. To correct for the stimulus-choice correlation that existed in the visual discrimination task (but not in the visually-independent auditory task, Supplementary Figure 4b), we subtracted from the stimulus-decoding accuracy at each timepoint a choice-decoding correction factor calculated as follows. We calculated the classification accuracy of the stimulus-trained decoder at predicting choice labels on difficult trials, using the same number of difficult trials as the stimulus test set, randomly drawn from the full set of difficult trials on each model repeat. Thus, the performance of the model that was due to actually decoding choice was removed by subtracting the mean accuracy of choice decoding on the correction set, leaving “true stimulus” decoding.

To assess whether the accuracy on the test set was significantly different from chance at a given timepoint, we trained a classifier on shuffled labels relative to the trial-by-trial stimulus activity. By repeating this on 100 shuffles of the data, we established a 95% confidence interval for each timepoint in each session. A classifier was labelled as significantly more accurate than chance if its test set accuracy exceeded the upper bound of the confidence interval. Comparisons to assess significance were done on a within-session basis to account for any structure arising from the distribution of trials on that session.

## Acknowledgments

We thank Priyanka Gupta, Fred Marbach, Torben Ott, and Petr Znamenskiy for comments on an earlier version of this manuscript. This work was supported by grant R01DC012565 from the National Institutes of Health (A.M.Z).

## Supplementary Figures

**Supplementary Figure 1.**
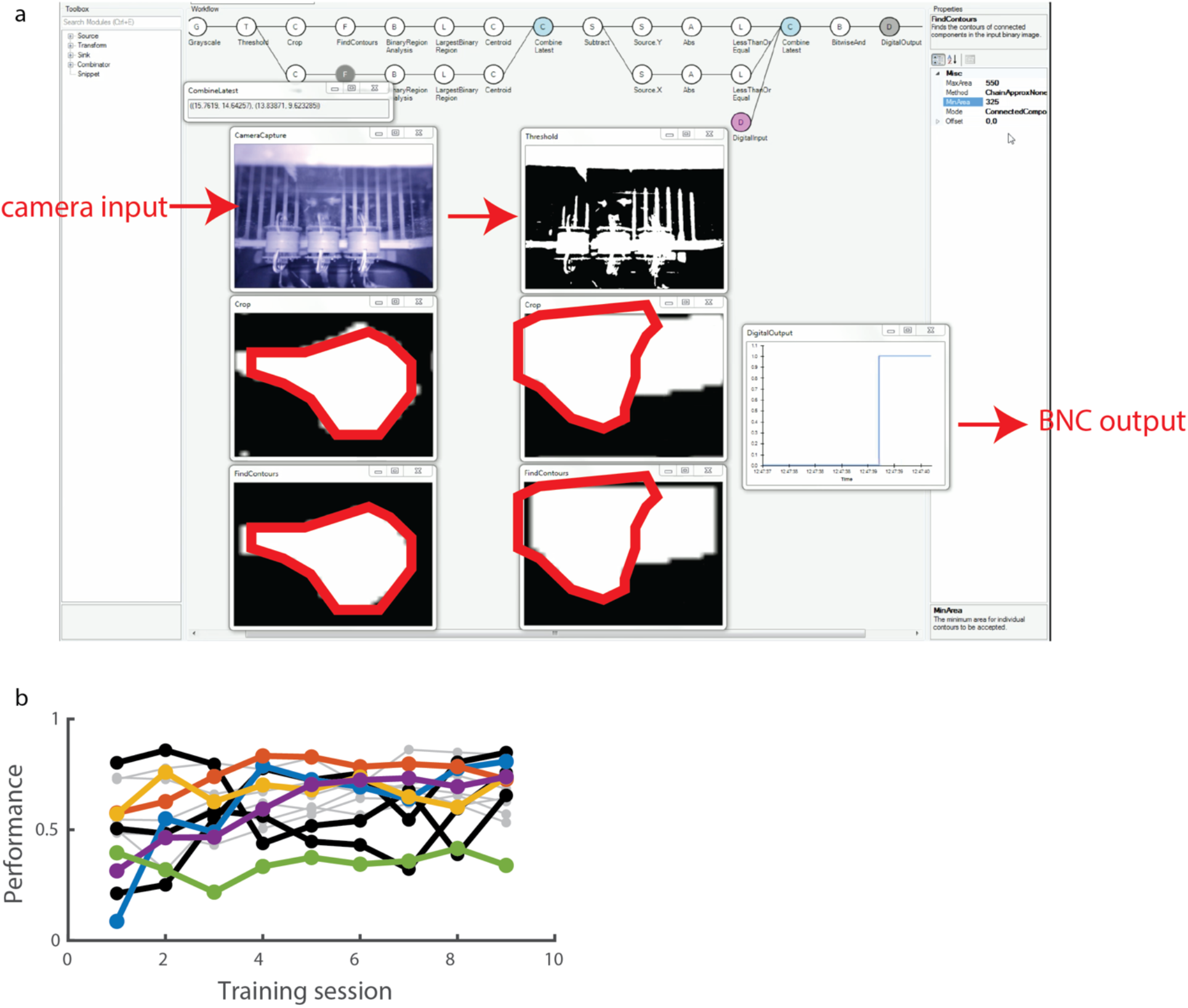
Bonsai-mediated online head position training. a. Bonsai workflow showing thresholding of camera input, identification of ROIs, and digital output to behavior control. Ear shapes are outlined in red for reference. b. Proportion of completed trials with training after introducing head position criterion.

**Supplementary Figure 2.**
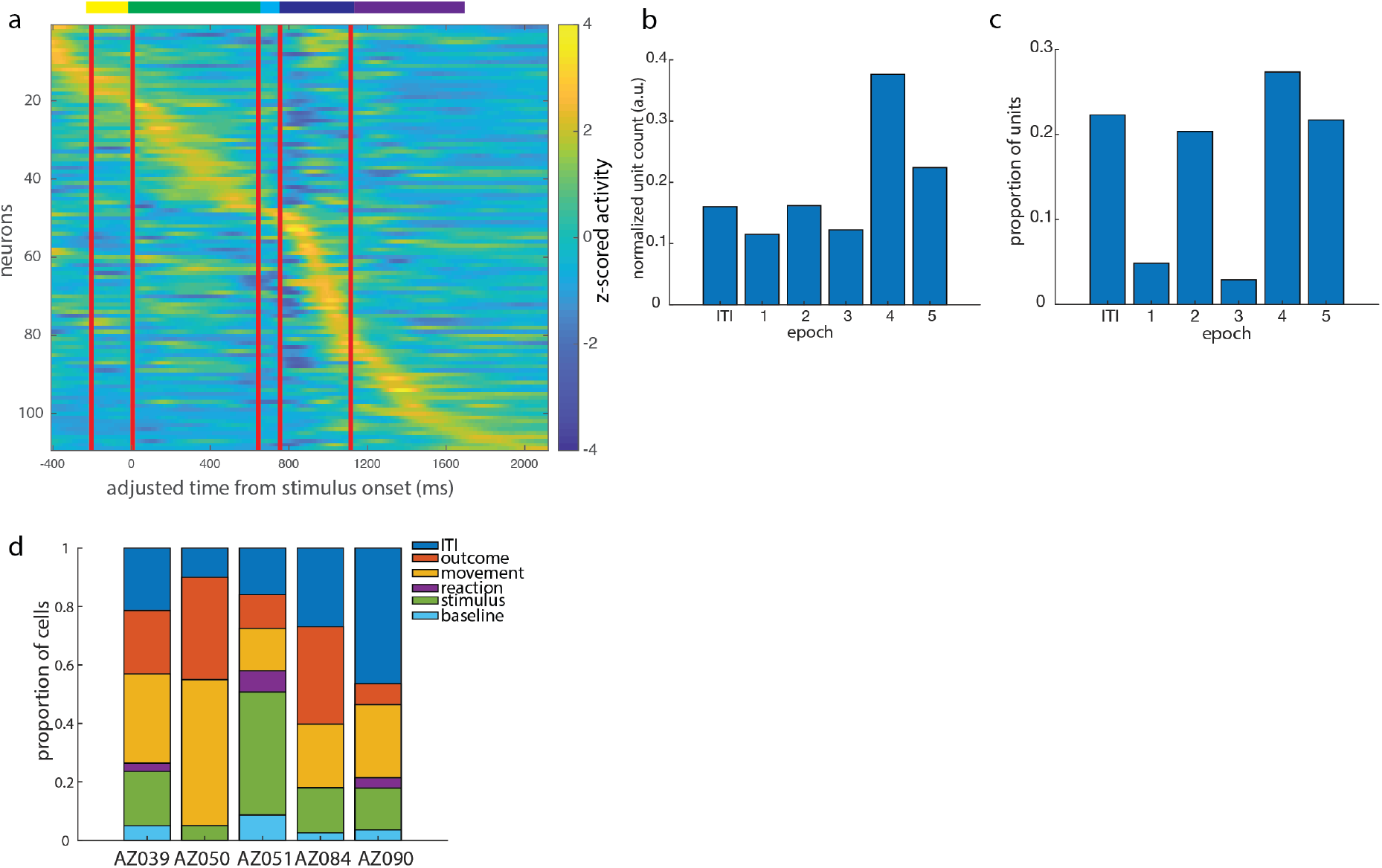
Timing of peak activity over recording dataset. a. Mean activity patterns of putative multi-units, sorted by peak activity timing. b. Counts of recorded units with peak in each epoch, normalized by epoch duration. c. Proportion of recorded units with peak in each epoch, as a proportion of recorded population. d. Peak activity timing distribution by animal.

**Supplementary Figure 3.**
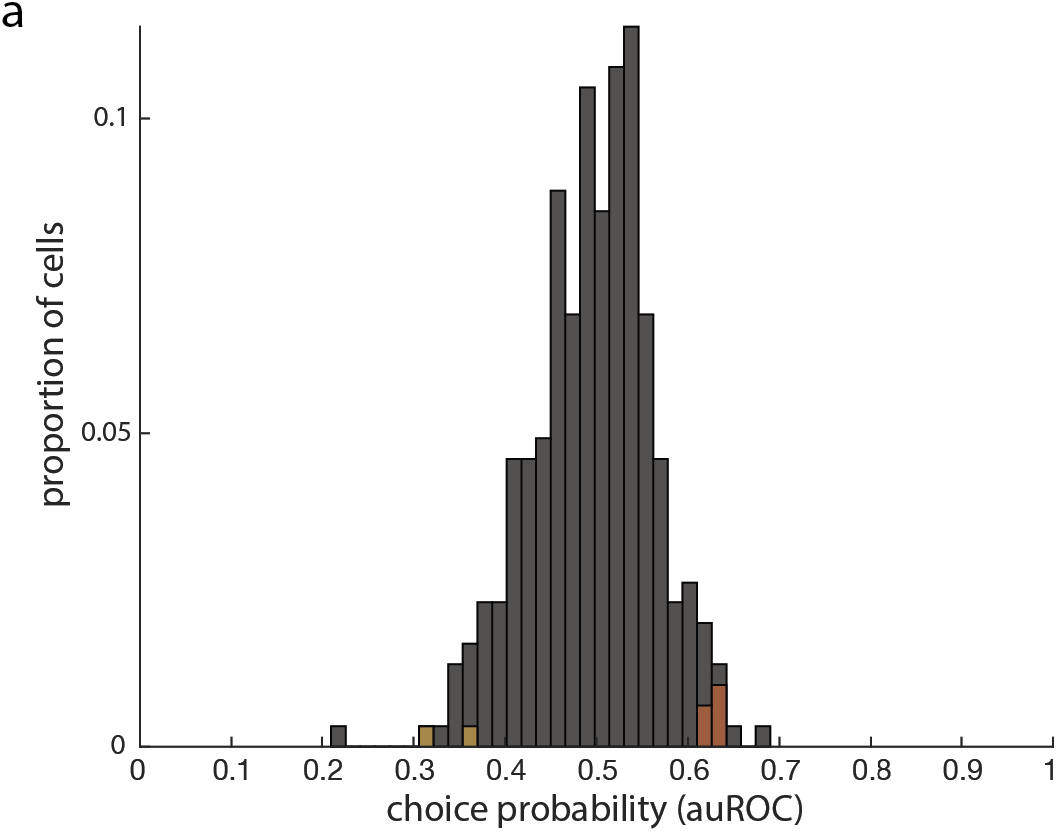
ROC analyses confirm low choice probabilities in V1.

**Supplementary Figure 4.**
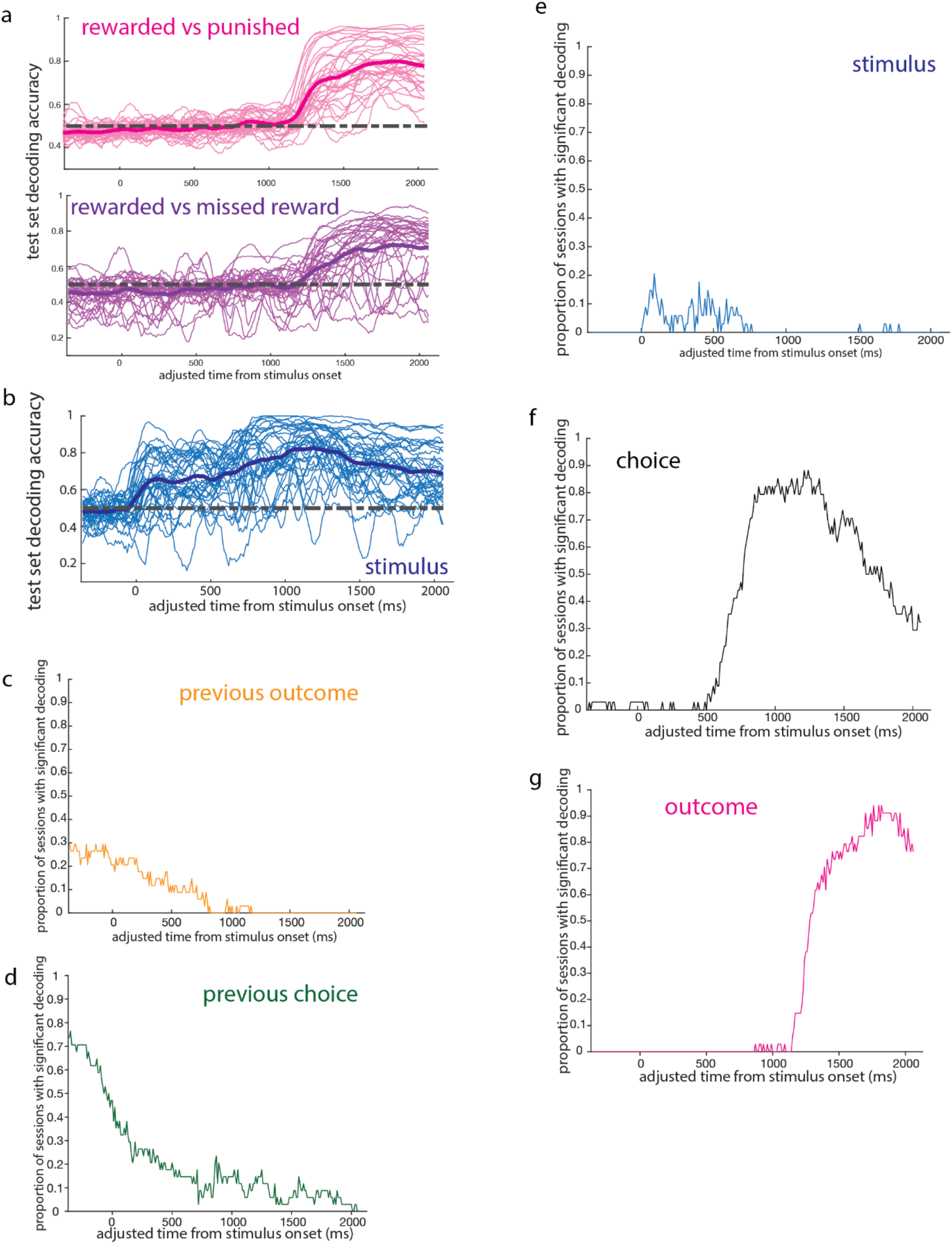
Significance testing of population decoding timecourses. a. Outcome epoch decoding is similar between decoding reward vs punishment, and reward vs missed reward. b. “Stimulus” decoding persists in choice epoch due to strong stimulus-choice correlation in trained animals. c-g. Proportion of sessions with decoding accuracy significantly above chance for each feature in Figure 6.

**Supplementary Figure 5.**
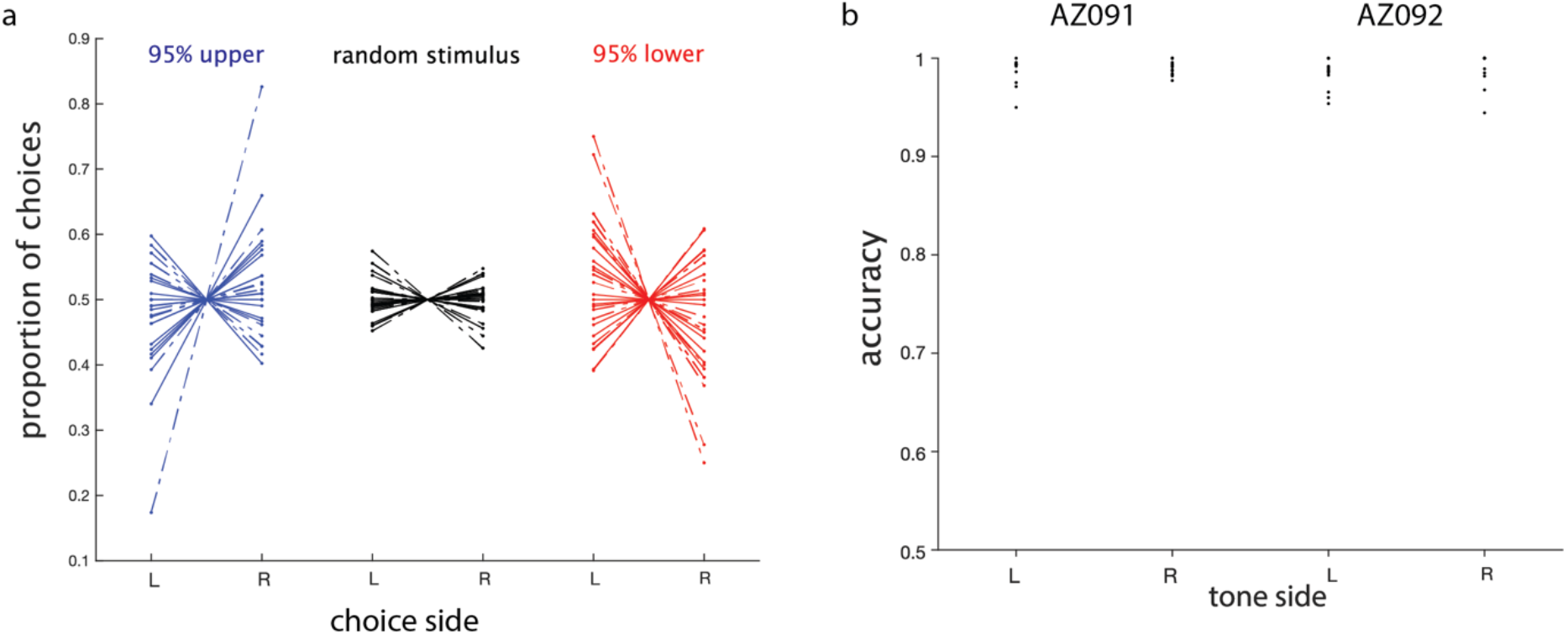
Behavior on a visually-independent decision task depends on tone location, not visual stimulus distribution. a. Proportion of left (L) and right (R) choices for both animals (AZ091: solid lines; AZ092: dashed lines) during each recording session, separated by visual stimulus identity. b. Decision accuracy, defined as choosing the same side as the go-tone was presented, remained stably above 90% across all recording sessions in both animals.

**Supplementary Figure 6.**
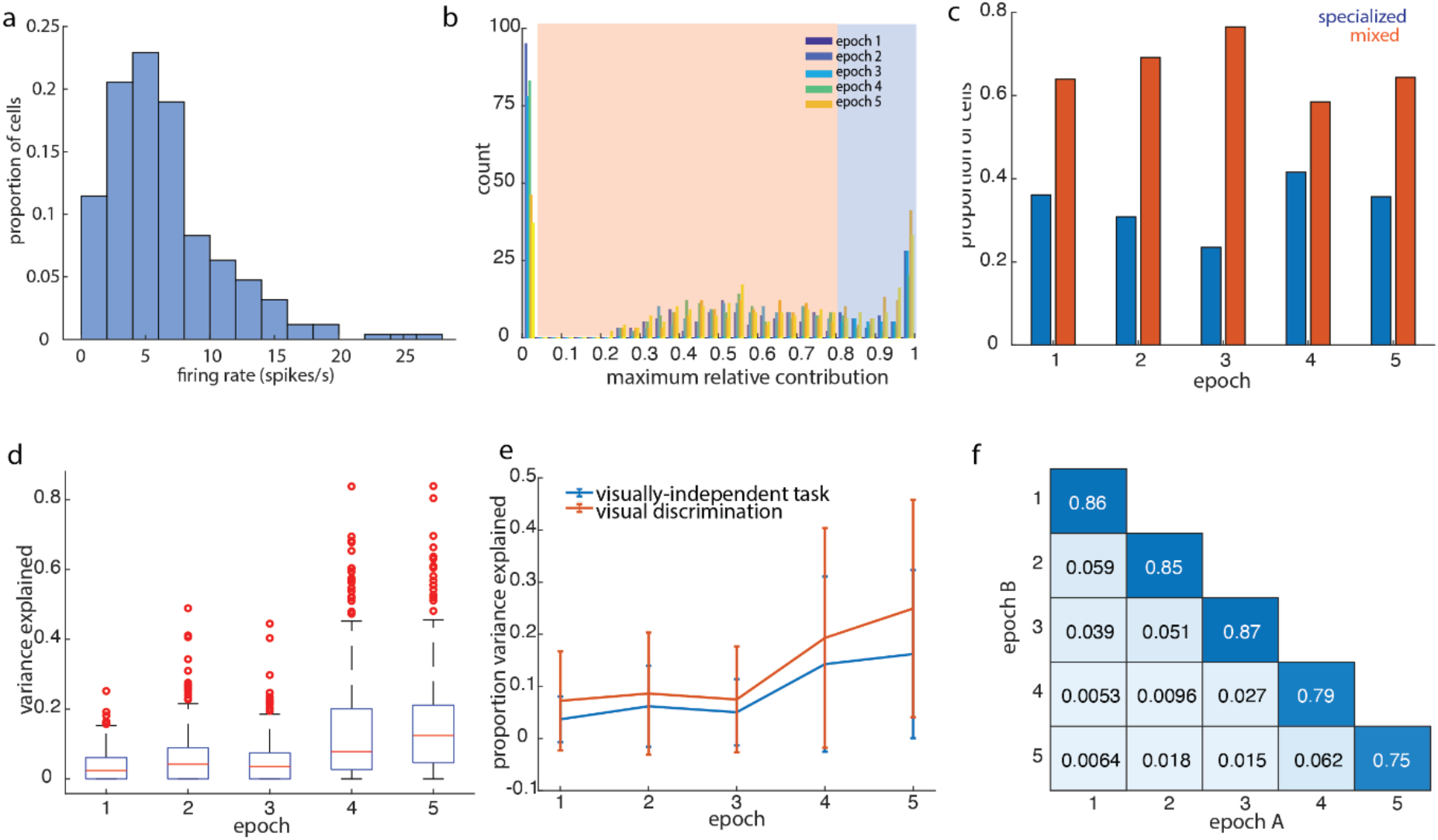
Linear encoding model reveals similar single neuron activity profiles between visual discrimination and visually-independent decision task. a. Firing rate distribution of single units recorded in visually-independent decision task. b. Distribution of maximum relative contribution of a single regressor to single neuron activity in the visually-independent decision task, by epoch. The same cutoff threshold separating “specialized” from “mixed” neurons as in the visual discrimination task is shown in shaded regions. c. Proportions of cells with “specialized” versus “mixed” selectivity profiles in the visually-independent task, as classified using the threshold in (a). d. Proportion of variance explained by linear encoding model in the visually-independent task, across behavioral epochs. e. Comparison of variance explained by linear model between visual discrimination task vs visually-independent decision task, across behavioral epochs. Points indicate mean, error bars indicate standard deviation. Median variance explained is significantly larger in the visual discrimination task than in the visually-independent decision task within each epoch (Mann-Whitney U-test, all p<0.005). f. Measure of cluster stability (adjusted Rand Index) when clustering single neuron feature encoding profiles between pairs of epochs, compared to stability over independent partitions in the same epoch (diagonal).

**Supplementary Figure 7.**
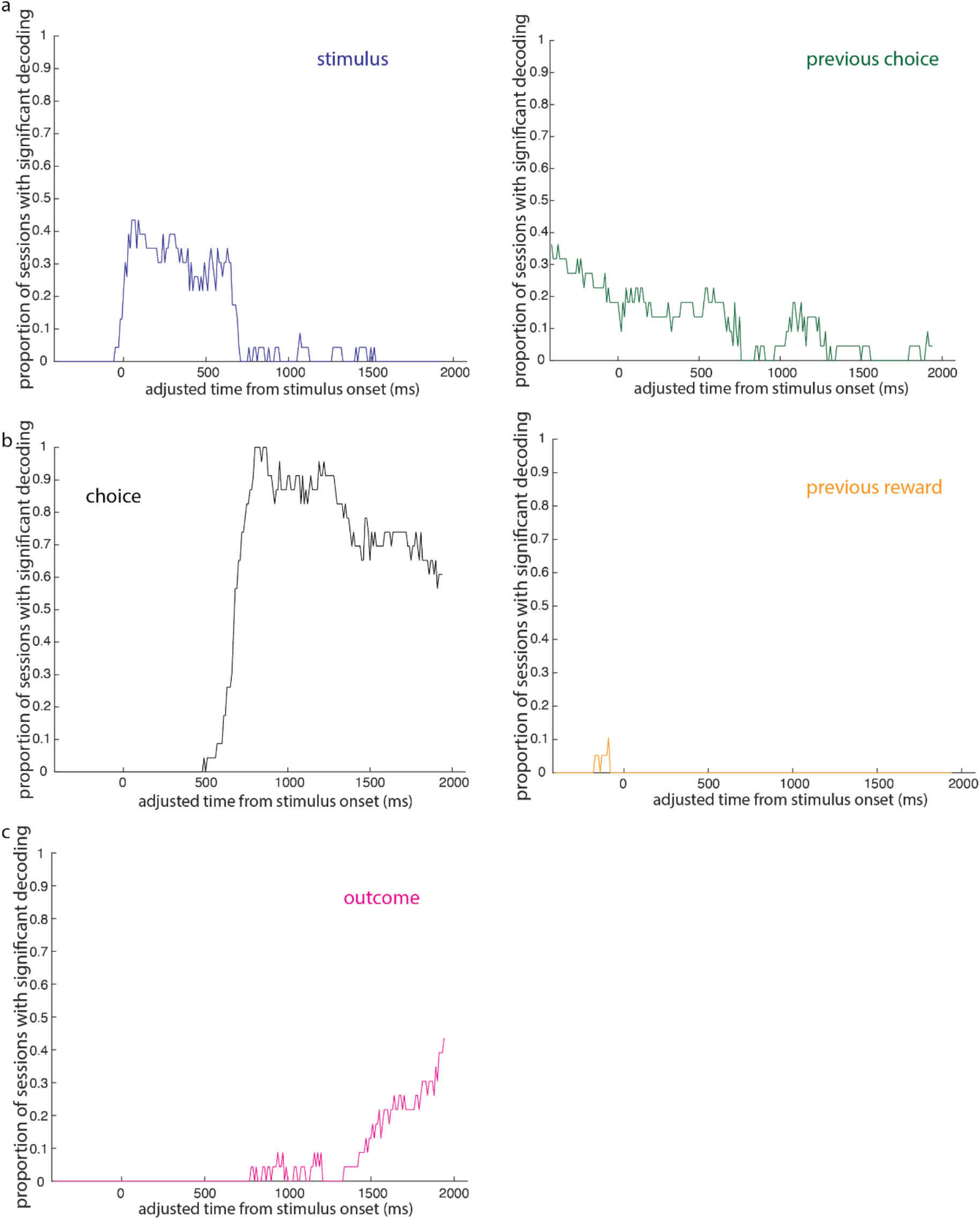
Significance testing of population decoding on visually-independent task.

## References

Brainard, D. H. (1997). The Psychophysics Toolbox. Spatial Vision 10(4): 433–36. https://doi.org/10.1163/156856897X00357

Engelhard, B., Finkelstein, J., Cox, J., Fleming, W., Jang, H. J., Ornelas, S., Koay, S. A., Thiberge, S. Y., Daw, N. D., Tank, D. W., & Witten, I. B. (2019). Specialized coding of sensory, motor and cognitive variables in VTA dopamine neurons. Nature, 570(7762), 509–513. https://doi.org/10.1038/s41586-019-1261-9

Felleman, D. J., & Van Essen D. C. (1991). Distributed Hierarchical Processing in the Primate Cerebral Cortex. Cerebral Cortex 1(1): 1–47. https://doi.org/10.1093/cercor/1.1.1-a

Fiser, A., Mahringer, D., Oyibo, H. K., Petersen, A. V., Leinweber, M., & Keller, G. B. (2016). Experience-dependent spatial expectations in mouse visual cortex. Nature neuroscience, 19(12), 1658–1664. https://doi.org/10.1038/nn.4385

Forys, B. J., Xiao, D., Gupta, P., & Murphy, T. H. (2020). Real-Time Selective Markerless Tracking of Forepaws of Head Fixed Mice Using Deep Neural Networks. eNeuro, 7(3), ENEURO.0096-20.2020. https://doi.org/10.1523/ENEURO.0096-20.2020

Guitchounts, G., Masís, J., Wolff, S., & Cox, D. (2020). Encoding of 3D Head Orienting Movements in the Primary Visual Cortex. Neuron, 108(3), 512–525.e4. https://doi.org/10.1016/j.neuron.2020.07.014

Hanks, T. D., Kopec, C. D., Brunton, B. W., Duan, C. A., Erlich, J. C., & Brody, C. D. (2015). Distinct relationships of parietal and prefrontal cortices to evidence accumulation. Nature, 520(7546), 220–223. https://doi.org/10.1038/nature14066

Hubel, D. H., Henson, C. O., Rupert, A., & Galambos, R. (1959). Attention units in the auditory cortex. Science, 129(3358), 1279–1280. https://doi.org/10.1126/science.129.3358.1279

Hubel, D. H., & Wiesel, T. N. (1959). Receptive fields of single neurones in the cat’s striate cortex. The Journal of physiology, 148(3), 574–591. https://doi.org/10.1113/jphysiol.1959.sp006308

Jaramillo, S., & Zador, A. M. (2011). The auditory cortex mediates the perceptual effects of acoustic temporal expectation. Nature neuroscience, 14(2), 246–251. https://doi.org/10.1038/nn.2688

Kane, G. A., Lopes, G., Saunders, J. L., Mathis, A., & Mathis, M. W. (2020). Real-time, low-latency closed-loop feedback using markerless posture tracking. eLife, 9, e61909. https://doi.org/10.7554/eLife.61909

Keller, G. B., Bonhoeffer, T., & Hübener, M. (2012). Sensorimotor mismatch signals in primary visual cortex of the behaving mouse. Neuron, 74(5), 809–815. https://doi.org/10.1016/j.neuron.2012.03.040

Keller, G. B., & Mrsic-Flogel, T. D. (2018). Predictive Processing: A Canonical Cortical Computation. Neuron, 100(2), 424–435. https://doi.org/10.1016/j.neuron.2018.10.003

Kleiner, M., Brainard, D., Pelli, D., Ingling, A., Murray, R., & Broussard, C. (2007). What’s new in psychtoolbox-3. Perception, 36(14), 1–16.

Krumin, M., Lee, J. J., Harris, K. D., & Carandini, M. (2018). Decision and navigation in mouse parietal cortex. eLife, 7, e42583. https://doi.org/10.7554/eLife.42583

Levy, S., Lavzin, M., Benisty, H., Ghanayim, A., Dubin, U., Achvat, S., Brosh, Z., Aeed, F., Mensh, B. D., Schiller, Y., Meir, R., Barak, O., Talmon, R., Hantman, A. W., & Schiller, J. (2020). Cell-Type-Specific Outcome Representation in the Primary Motor Cortex. Neuron, 107(5), 954–971.e9. https://doi.org/10.1016/j.neuron.2020.06.006

Lopes, G., Bonacchi, N., Frazão, J., Neto, J. P., Atallah, B. V., Soares, S., Moreira, L., Matias, S., Itskov, P. M., Correia, P. A., Medina, R. E., Calcaterra, L., Dreosti, E., Paton, J. J., & Kampff, A. R. (2015). Bonsai: an event-based framework for processing and controlling data streams. Frontiers in neuroinformatics, 9, 7. https://doi.org/10.3389/fninf.2015.00007

Marques, T., Summers, M. T., Fioreze, G., Fridman, M., Dias, R. F., Feller, M. B., & Petreanu, L. (2018). A Role for Mouse Primary Visual Cortex in Motion Perception. Current biology: CB, 28(11), 1703–1713.e6. https://doi.org/10.1016/j.cub.2018.04.012

Mathis, A., Mamidanna, P., Cury, K. M., Abe, T., Murthy, V. N., Mathis, M. W., & Bethge, M. (2018). DeepLabCut: markerless pose estimation of user-defined body parts with deep learning. Nature neuroscience, 21(9), 1281–1289. https://doi.org/10.1038/s41593-018-0209-y

Morcos, A. S., & Harvey, C. D. (2016). History-dependent variability in population dynamics during evidence accumulation in cortex. Nature neuroscience, 19(12), 1672–1681. https://doi.org/10.1038/nn.4403

Musall, S., Kaufman, M. T., Juavinett, A. L., Gluf, S., & Churchland, A. K. (2019). Single-trial neural dynamics are dominated by richly varied movements. Nature neuroscience, 22(10), 1677–1686. https://doi.org/10.1038/s41593-019-0502-4

Niell, C. M., & Stryker, M. P. (2008). Highly selective receptive fields in mouse visual cortex. The Journal of neuroscience: the official journal of the Society for Neuroscience, 28(30), 7520–7536. https://doi.org/10.1523/JNEUROSCI.0623-08.2008

Niell, C. M., & Stryker, M. P. (2010). Modulation of visual responses by behavioral state in mouse visual cortex. Neuron, 65(4), 472–479. https://doi.org/10.1016/j.neuron.2010.01.033

Nienborg, H., & Cumming, B. G. (2006). Macaque V2 neurons, but not V1 neurons, show choice-related activity. The Journal of neuroscience: the official journal of the Society for Neuroscience, 26(37), 9567–9578. https://doi.org/10.1523/JNEUROSCI.2256-06.2006

Orsolic, I., Rio, M., Mrsic-Flogel, T. D., & Znamenskiy, P. (2021). Mesoscale cortical dynamics reflect the interaction of sensory evidence and temporal expectation during perceptual decision-making. Neuron, 109(11), 1861–1875.e10. https://doi.org/10.1016/j.neuron.2021.03.031

Osako, Y., Ohnuki, T., Tanisumi, Y., Shiotani, K., Manabe, H., Sakurai, Y., & Hirokawa, J. (2021). Contribution of non-sensory neurons in visual cortical areas to visually guided decisions in the rat. Current biology: CB, 31(13), 2757–2769.e6. https://doi.org/10.1016/j.cub.2021.03.099

Otazu, G. H., Tai, L. H., Yang, Y., & Zador, A. M. (2009). Engaging in an auditory task suppresses responses in auditory cortex. Nature neuroscience, 12(5), 646–654. https://doi.org/10.1038/nn.2306

Paxinos, G., & Watson, C. (2007). The rat brain in stereotaxic coordinates. Amsterdam: Elsevier.

Pelli, DG (1997). The VideoToolbox software for visual psychophysics: Transforming numbers into movies, Spatial Vision 10(4):437–442. https://doi.org/10.1163/156856897X00366

Poort, J., Khan, A. G., Pachitariu, M., Nemri, A., Orsolic, I., Krupic, J., Bauza, M., Sahani, M., Keller, G. B., Mrsic-Flogel, T. D., & Hofer, S. B. (2015). Learning Enhances Sensory and Multiple Non-sensory Representations in Primary Visual Cortex. Neuron, 86(6), 1478–1490. https://doi.org/10.1016/j.neuron.2015.05.037

Puścian, A., Benisty, H., & Higley, M. J. (2020). NMDAR-Dependent Emergence of Behavioral Representation in Primary Visual Cortex. Cell reports, 32(4), 107970. https://doi.org/10.1016/j.celrep.2020.107970

Raposo, D., Kaufman, M. T., & Churchland, A. K. (2014). A category-free neural population supports evolving demands during decision-making. Nature neuroscience, 17(12), 1784–1792. https://doi.org/10.1038/nn.3865

Rigotti, M., Barak, O., Warden, M. R., Wang, X. J., Daw, N. D., Miller, E. K., & Fusi, S. (2013). The importance of mixed selectivity in complex cognitive tasks. Nature, 497(7451), 585–590. https://doi.org/10.1038/nature12160

Saleem, A. B., Diamanti, E. M., Fournier, J., Harris, K. D., & Carandini, M. (2018). Coherent encoding of subjective spatial position in visual cortex and hippocampus. Nature, 562(7725), 124–127. https://doi.org/10.1038/s41586-018-0516-1

Scott, B. B., Constantinople, C. M., Akrami, A., Hanks, T. D., Brody, C. D., & Tank, D. W. (2017). Fronto-parietal Cortical Circuits Encode Accumulated Evidence with a Diversity of Timescales. Neuron, 95(2), 385–398.e5. https://doi.org/10.1016/j.neuron.2017.06.013

Scott, B. B., Brody, C. D., & Tank, D. W. (2013). Cellular resolution functional imaging in behaving rats using voluntary head restraint. Neuron, 80(2), 371–384. https://doi.org/10.1016/j.neuron.2013.08.002

Shuler, M. G., & Bear, M. F. (2006). Reward timing in the primary visual cortex. Science (New York, N.Y.), 311(5767), 1606–1609. https://doi.org/10.1126/science.1123513

Steinmetz, N. A., Zatka-Haas, P., Carandini, M., & Harris, K. D. (2019). Distributed coding of choice, action and engagement across the mouse brain. Nature, 576(7786), 266–273. https://doi.org/10.1038/s41586-019-1787-x

Stringer, C., Pachitariu, M., Steinmetz, N., Reddy, C. B., Carandini, M., & Harris, K. D. (2019). Spontaneous behaviors drive multidimensional, brainwide activity. Science (New York, N.Y.), 364(6437), 255. https://doi.org/10.1126/science.aav7893

Uchida, N., & Mainen, Z. F. (2003). Speed and accuracy of olfactory discrimination in the rat. Nature neuroscience, 6(11), 1224–1229. https://doi.org/10.1038/nn1142

Vinck, M., Batista-Brito, R., Knoblich, U., & Cardin, J. A. (2015). Arousal and locomotion make distinct contributions to cortical activity patterns and visual encoding. Neuron, 86(3), 740–754. https://doi.org/10.1016/j.neuron.2015.03.028

Wallace, D. J., Greenberg, D. S., Sawinski, J., Rulla, S., Notaro, G., & Kerr, J. N. (2013). Rats maintain an overhead binocular field at the expense of constant fusion. Nature, 498(7452), 65–69. https://doi.org/10.1038/nature12153

